# Direct observation of assembly and function of trigger responsive lipase biohybrids

**DOI:** 10.1101/2024.12.20.629668

**Authors:** Errika Voutyritsa, Emily Winther Sørensen, Athanasios Oikonomou, Thomai Lazou, Jonida Bushi, Sune M. Christensen, Kelly Velonia, Nikos S. Hatzakis

## Abstract

Protein-polymer biohybrids mark a cutting edge in the creation of advanced materials designed for high performance under diverse conditions by combining the intrinsic stability and adaptability of synthetic polymers with the remarkable functionality and specificity of proteins. Despite the existence of stimuli responsive polymers, their potential for achieving precise, trigger responsive control over both the function and the assembly of protein-polymer biohybrids has yet to be fully attained. Here, we report the synthesis and thorough characterization of pH and temperature responsive lipase-polymer conjugates at both the supramolecular and the single molecule level. In contrast to conventional ensemble measurements performed in solution, we extensively characterized the stimuli triggered reversibility of the self-assembly of these lipase-based biohybrids and recorded the enzymatic activity at the fundamental level of individual nanoparticles. Our findings unexpectedly demonstrated that the stimuli responsive biohybrids exhibited not only a twofold increased enzymatic activity as compared to native lipases in certain cases but also maintained the trigger responsive control over enzymatic activity by temperature. Direct single particle recordings of biohybrid activity showed a correlation of the biohybrid size to catalytic activity, revealing individual enzymes to display higher activity in smaller nanoparticles. The biohybrids exhibit impressive long-term stability, in some cases almost 95% of the enzymatic activity as compared to ∼40-60% for native lipases after 1 year of storage. The improved stability and activity as compared to native enzymes accompanied with their triggered responsiveness highlight their significant potential for tightly controlled biocatalysis and sustainable industrial applications.

## INTRODUCTION

Protein-polymer biohybrids have emerged as a transformative class of materials, uniquely combining the biocompatibility, specificity, and functionality of proteins with the versatility and structural stability of synthetic polymers.^1-11^ Their potential has been demonstrated across biomedical, industrial, and environmental applications, such as drug delivery,^12-15^ and catalysis.^8,16-19^ Depending on the protein and polymer component amphiphilic protein-polymer conjugates form assemblies with diverse morphologies from vesicular structures to defined nanoparticles and dimensions ranging from nanometres to micrometres.^20^

Notable, focus has been directed towards the development of responsive protein-polymer biohybrids capable of altering their physicochemical properties and function in response to environmental stimuli such as pH, temperature, and ionic strength.^21-28^ Temperature-sensitive biohybrids formulated using polymers like poly(*N*-isopropylacrylamide) (PNIPAM),^21-23,28^ that undergoes temperature dependent phase transitions, allow for temperature-mediated control of their properties. Design of pH-responsive biohybrids using polyacrylic acids^29^ or polyamino acrylates,^30^ offer the possibility of pH specific responses ideal for targeted drug delivery in acidic tumor microenvironments or intracellular compartments. While these stimuli-responsive properties unlock the potential of biohybrids to perform dynamically and respond to environmental cues, this potential is yet to be fully realised and understood. Current performance of such protein-polymer biohybrids in biocatalysis,^8,16-19^ which is investigated in not trigger responsive systems, and drug delivery^12-15^ primarily rely on bulk measurements, masking system heterogeneity and averaging the interplay between dimensions, and functional output.

Here, pH and temperature-responsive polymers were grafted from *Candida antarctica Lipase B* (CALB) and *Thermomyces lanuginosus Lipase* (TLL) and produced stimuli responsive biohybrids that can serve as trigger responsive biocatalysts, that structurally and functionally respond to specific environmental cues (Fig. **1A**). Besides exhibiting enhanced catalytic activity in some cases, outperforming the native enzymes counterparts, the biohybrids displayed increased thermal stability and long-term storage. Moreover, the responsiveness of the polymeric moiety enables modulation and control of the catalytic activity of the biohybrids under varying pH and temperature conditions. Thorough characterisation of individual nanoparticles at the fundamental level revealed stark heterogeneities in their growth rates (number of unimers assembling over time), how those correlates to nanoparticle dimensions, as well as a hitherto unknown correlation of the dimensions of the self-assembled biohybrids assemblies to the function of each individual enzyme. The obtained single particle readouts surprisingly revealed that enzymes incorporated in smaller assemblies display higher activity than those integrated in larger assemblies. Besides enhancing our understanding on structure-function relationships within biohybrid systems, these findings pave the way for the design and development of smart, efficient, and trigger responsive functional biomaterials for a host of biomedical and biocatalytic applications.

**Figure 1:**
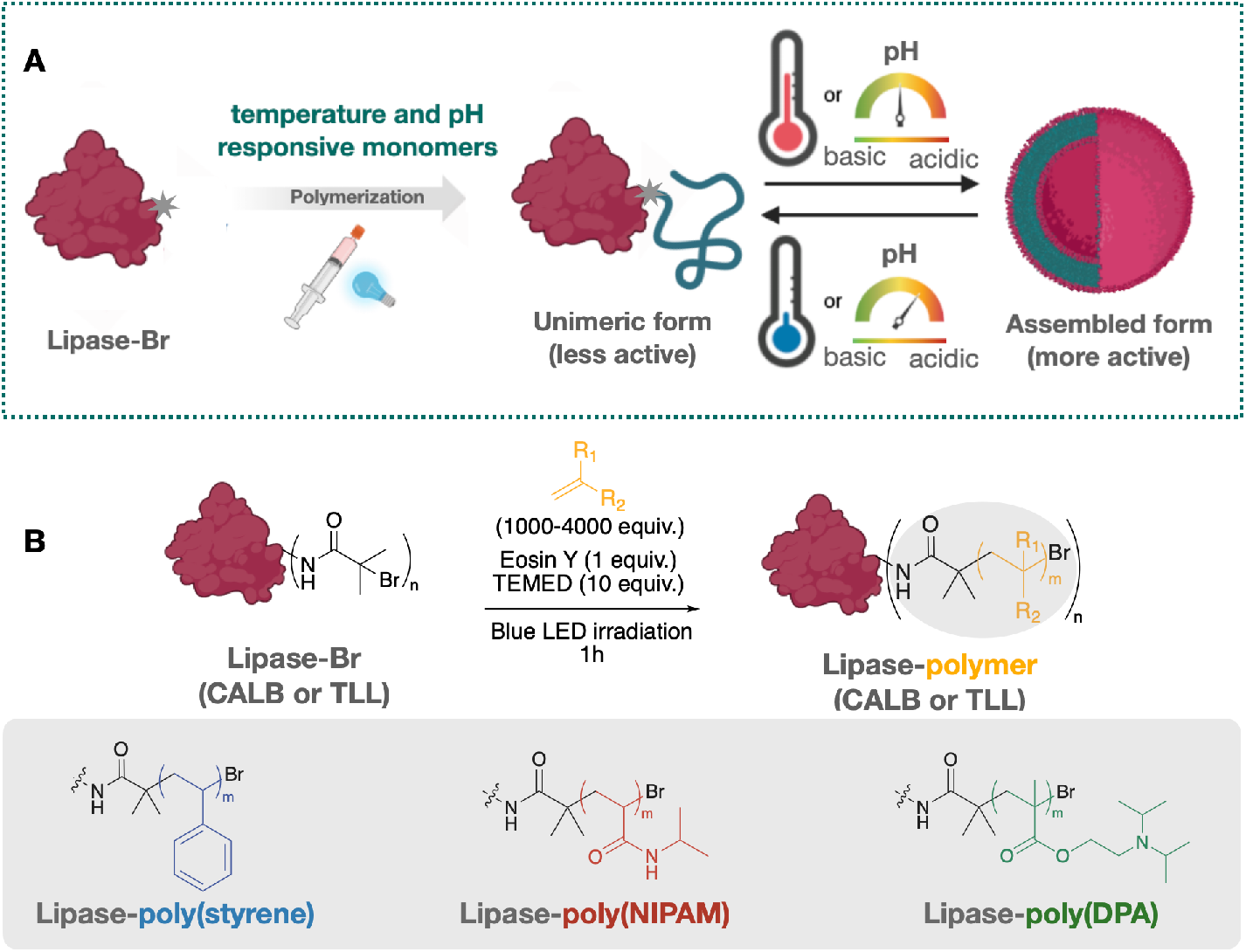
Synthesis and characterisation of CALB and TLL biohybrids. **A**. Schematic representation of the temperature and the pH triggered response of temperature and pH responsive lipase based biohybrids. **B**. Schematic representation of the Eosin Y photocatalyzed synthesis of the biohybrids.

## RESULTS AND DISCUSSION

### Single Particle Characterization and Reversibility of Responsive Enzyme Biohybrids

The responsive biohybrids were synthesised following an Eosin Y photocatalyzed oxygen tolerant grafting from approach (Fig. **1B**).^27^ As model biocatalysts we employed CALB and TLL that together with other lipases constitute major class of enzymes used as biocatalysts.^31-35^ Styrene, *N*-isopropylacrylamide (NIPAM), and 2-(diisopropylamino)ethyl methacrylate (DPA) were selected as the monomers. Synthesis was performed under oxygen tolerant conditions, following a protocol recently reported by us^27^ (Fig. **1B**, see methods and Supplementary Information).

The response of the temperature responsive biohybrids CALB- and TLL-poly(NIPAM) was initially evaluated by DLS measurements below and above the Low Critical Solution Temperature of 32 °C. At 25 °C the hydrodynamic diameter of CALB- and TLL-poly(NIPAM) were found approximately 13.50 nm and 14.14 nm respectively, suggesting that they exist in a monomeric state. At 37 °C, the diameters increased to 327.4 nm for CALB-poly(NIPAM) and 163.7 nm for TLL-poly(NIPAM), confirming the onset of the assembly process (Fig. **2E**, **S10A** and **S16A**). Interestingly, kinetic measurements for prolonged periods at 37 °C showed that TLL-poly(NIPAM) reaches equilibrium faster that CALB-poly(NIPAM) (Fig. **S12, S18**). Further, a systematic variation of temperature from 20 °C to 45 °C revealed a continuous increase in the average diameter of CALB-poly(NIPAM) and a plateau in the diameter increase of TLL-poly(NIPAM) beyond 30 °C (Fig. **S11, S17**,Table **S2, S7)**. This difference in kinetics may originate from the higher hydrophobicity of TLL.^36^

**Figure 2.**
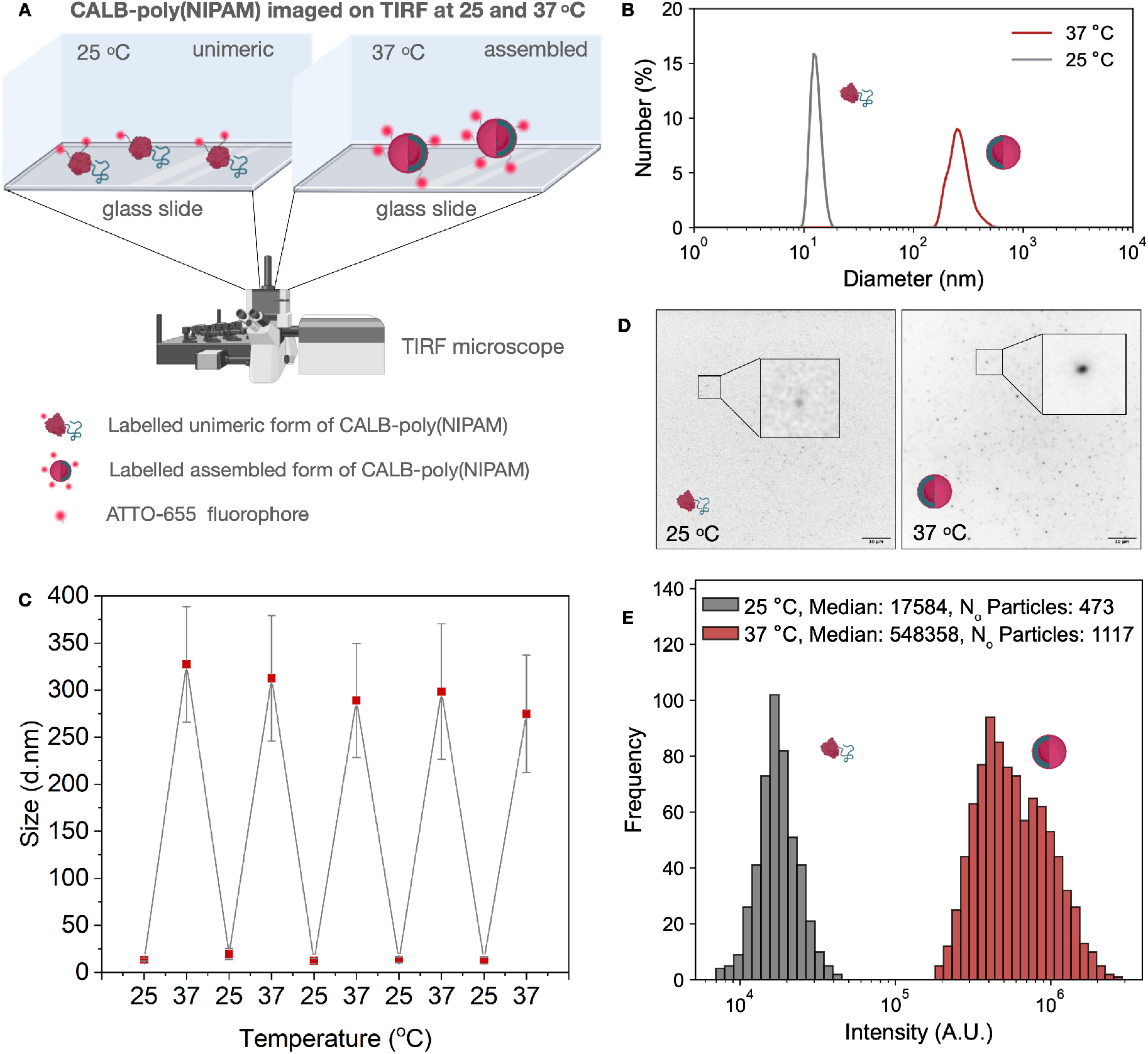
Characterization of the temperature-dependent reversible assembly of spherical CALB-poly(NIPAM) structures. **A**. Schematic representation of the experimental setup for direct observation of ATTO-655 labelled temperature responsive CALB-poly(NIPAM) at two different temperatures on a TIRF microscope in two independent experiments. At 25 °C CALB-poly(NIPAM) was found in a unimeric form, whereas at 37 °C CALB-poly(NIPAM) is a self-assembled form resulting in increased signal. **B**. Overlay of DLS measurements displaying the hydrodynamic diameter distribution of CALB-poly(NIPAM) at 25 °C and 37 °C. At 25 °C the mean hydrodynamic diameter was 13.50 +/-1.56 with a PDI of 11.2. At 37 °C the mean hydrodynamic diameter was 327.40 +/- 61.44 with a PDI of 19.2. Error bars correspond to the standard deviation of the size distribution. **C**. Reversibility of CALB-poly(NIPAM) temperature-induced response as characterized by DLS reporting the average hydrodynamic diameter of CALB-poly(NIPAM) upon undergoing five cycles of temperature transitions from 25 °C to 37 °C. Error bars correspond to the standard deviation of the size distribution. **D**. Representative experimentally TIRF images of CALB-poly(NIPAM) at 25 °C and 37 °C displaying the biohybrid at its unimeric and assembled form respectively. **E**. Background corrected intensity distributions of CALB-poly(NIPAM) labelled with ATTO-655 at 25 °C and 37 °C. At 25 °C the intensity distribution has median of 17583.6 with a sigma of 4348.0. The number of detected particles is 473. At 37 °C the intensity distribution has median of 548358.1 with a sigma of 256153.5. The number of detected particles is 1117. Intensity axis is scaled logarithmically. CALB-poly(NIPAM) was imaged at 25 °C and 37 °C as two independent experiments. Data correspond to intensities imaging at 6 fields of view for each experiment.

To test the reversibility of the assembly process, we measured by DLS the hydrodynamic radius upon repeated temperature cycling between 25 °C and 37 °C at which temperature TLL-poly(NIPAM) was shown to reach its size plateau while CALB-poly(NIPAM) was in a self-assembled morphology, although without having reached an equilibrium. The consistent changes in the hydrodynamic diameter observed for both temperature responsive biohybrids demonstrated their robustness and reliable temperature sensitivity (Fig. **2F**, **S10B** and **S16B**, Tables **S1** and **S6**).

To quantify the heterogeneity of the diameter of the CALB- and TLL-poly(NIPAM) assemblies we employed single particle fluorescence readouts.^33,34,37-45^ We labelled the enzyme component of CALB- and TLL-poly(NIPAM) using ATTO-655 NHS and extracted their dimension using TIRF microscopy under conditions favouring either the monomeric or the assembled forms. To accurately assess the size of the diffraction limited spots and its dependence on the response conditions, we relied on our recent work.^40^ The single particle readout allowed simultaneous recordings of hundreds of fluorescently labelled biohybrids revealing significant temperature-dependent size increase (Fig. **2**) (*see methods and supplementary information*).^40^ The intensity of CALB-poly(NIPAM) demonstrated a 30-fold increase from 25 °C to 37 °C, whereas the intensity of TLL-poly(NIPAM) showed a 15-fold increase under the same temperature variation, signifying the assembly process leading to the formation of larger nanoparticles (Fig. **2C** and **2D**, **S21**). To observe directly the disassembly process of the assembled of CALB- and TLL-poly(NIPAM) biohybrids, we decreased the temperature from 37 °C to 25 °C, which resulted in temperature triggered changes in particle count and morphology. An 8-fold decrease in intensity was observed for CALB-poly(NIPAM) with similar trends seen for TLL-poly(NIPAM) (Fig. **S28**), highlighting the reversibility of the nanoparticle formation.

We then explored the pH responsiveness of CALB- and TLL-poly(DPA). DLS showed the existence of the unimer and/or dimer at pH 4.0 and of the assembled particles even at pH as low as 5.0. The dimensions of the nanoparticles progressively increased upon raising the pH to 8.0 (Fig. **3D**). Consistent with DLS, single particle measurements showed a wide distribution of diameters at pH 8.2 with sizes ranging from 30 to 150 nm for CALB-poly(DPA) (Fig. **3B**), whereas similar results were obtained for TLL-poly(DPA) (Fig. **3C**, **S22** and **S25**), highlighting the size distribution heterogeneity of the biohybrids.

**Figure 3.**
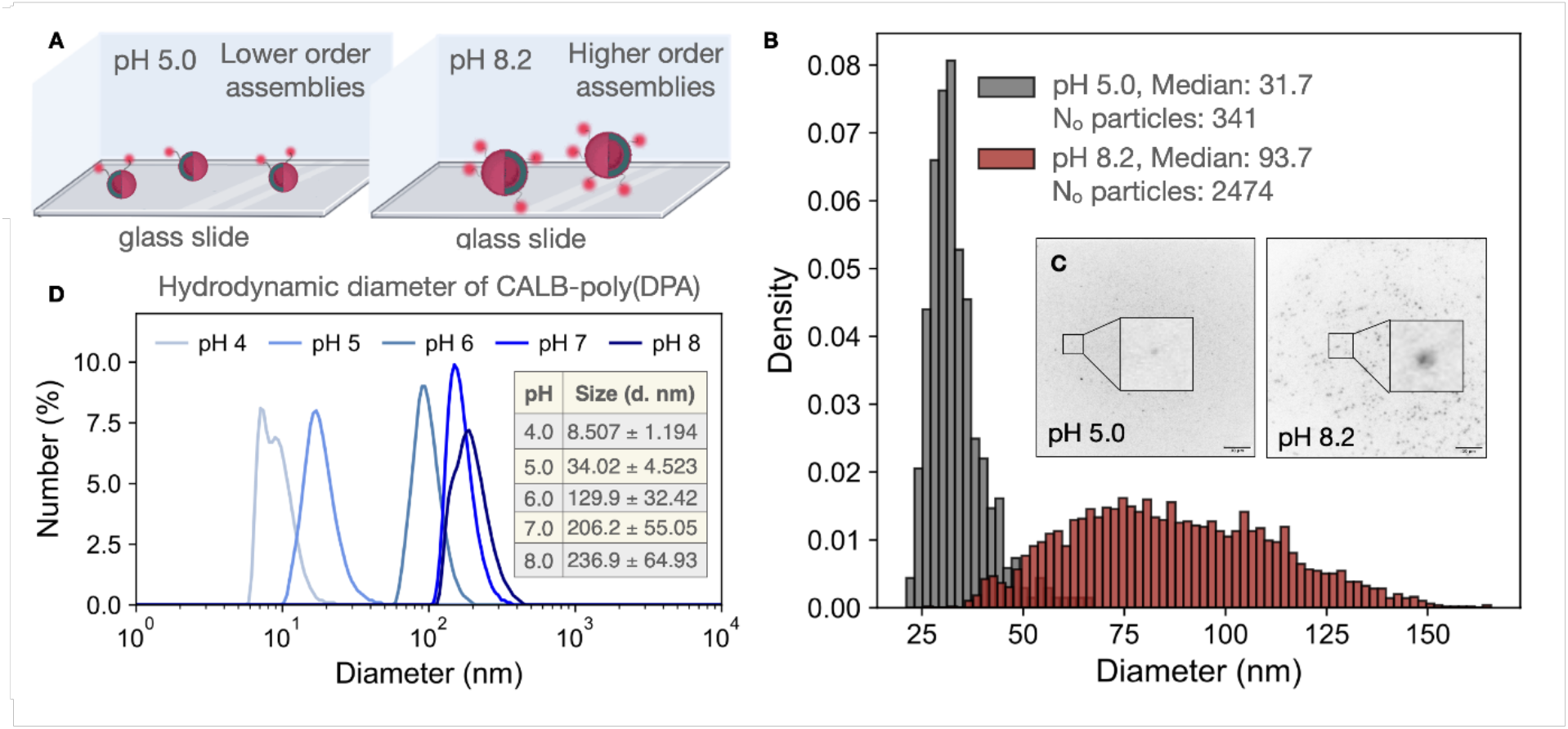
Characterisation of the pH dependent response and assembly of higher order vesicular structures of CALB-poly(DPA). **A**. Cartoon representation of the experimental setup for direct observation of ATTO-655 labelled pH responsive CALB-poly(DPA) at 2 different pH values on TIRF microscope as 2 independent experiments. At pH 5.0 CALB-poly(DPA) forms lower order nanoparticles, whereas at pH 8.2 CALB-poly(DPA) forms higher order nanoparticles. **B**. Size distributions CALB-poly(DPA) at pH 5.0 and 8.2. Diameter values are extracted as previously mentioned (see methods).^44^ At pH 5.0 the size distribution has median of 31.742 with a sigma of 4.683. The number of detected particles is 341. At pH 8.2 the size distribution has median of 93.689 with a sigma of 34.785. The number of detected particles is 2474. Inset: **C**. Representative experimentally acquired TIRF images of CALB-poly(DPA) at pH 5.0 and pH 8.2 displaying the biohybrid forming lower and higher order nanoparticles respectively **D**. pH dependence of the hydrodynamic diameter distribution of CALB-poly(DPA) at pH 4.0, 5.0, 6.0, 7.0 and 8.0 measured on DLS. Data corresponds to DLS overlays recorded at pH 4.0, 5.0, 6.0, 7.0 and 8.0. All the samples were incubated at each pH value for 24 hours before measurement. Mean values correspond to the mean hydrodynamic diameter at each pH value. Error bars correspond to the standard deviation of the size distribution.

Quantitative image analysis enabled the extraction of the number of enzymes per assembled nanoparticle using the equation (i)^40^ where DOL corresponds to Degree of Labelling (Tables 1 and S10). Poly(styrene) and poly(DPA) biohybrids, already assembled under labelling conditions, capture only surface exposed enzymes. In contrast, poly(NIPAM) biohybrids were labelled prior assembly anticipating all enzymes to be labelled. As expected, at 25 °C, CALB- and TLL-poly(NIPAM) exist in the form of unimers displaying similar intensity with single enzymes (see Fig. S26), whereas at 37 °C these biohybrids form nanoparticles consisting of ∼11k enzymes. Notably, at 37°C, CALB-poly(NIPAM) forms assemblies with diameters of ∼340 nm, while TLL-poly(NIPAM) forms significantly smaller assemblies with diameters of ∼60 nm (Table 1, entries 3 and 8). Despite this size difference, both biohybrids contain the same number of enzymes, suggesting a denser, more tightly packed enzyme arrangement in the TLL-poly(NIPAM) biohybrids. Along the same lines, CALB-poly(DPA) forms assemblies with varying sizes depending on the pH with the number of enzymes to vary from 559 to ∼6k from pH 5 to pH 8 respectively. Similarly, TLL-poly(DPA) forms smaller assemblies of ∼1K enzymes at pH 5 and larger of ∼11k enzymes at pH 8. For both CALB- and TLL-poly(DPA), the enzyme count aligns with the nanoparticle size, indicating consistent polymer-driven packing density (Table 1, entries 4,5,9,10). Interestingly, the assembled form of TLL-poly(NIPAM) and TLL-poly(DPA) at pH 8 contains a significantly higher number of enzymes than expected, suggesting the formation of a multilayer assembly (Table 1, entries 8,9).

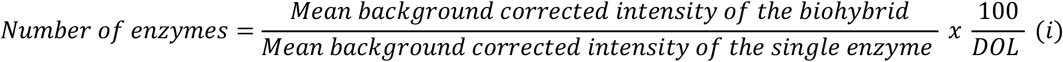

**Table 1.** Calculation of the mean number of the enzymes per biohybrid assembly. Data corresponds to 6 technical replicates. Error bars correspond to the standard deviation of the mean.

| Entry | Biohybrid | Temperature (°C) | pH | Mean diameter (DLS) | N<sub>o</sub> of enzymes |
| --- | --- | --- | --- | --- | --- |
| 1 | CALB coated poly(styrene) | 25 | 8.2 | 83.55 | 2145.657 +/- 362.616 |
| 2 | CALB-poly(NIPAM) | 25 | 8.2 | 11.54 | Unimer |
| 3 | CALB-poly(NIPAM) | 37 | 8.2 | 345.3 | 11029.844 +/- 4870.413 |
| 4 | CALB-poly(DPA) | 25 | 5.0 | 31.69 | 559.053 +/- 132.504 |
| 5 | CALB-poly(DPA) | 25 | 8.2 | 87.71 | 5940.486 +/- 1407.989 |
| 6 | TLL-poly(styrene) | 25 | 8.2 | 89.27 | 2601.306 +/- 530.795 |
| 7 | TLL-poly(NIPAM) | 25 | 8.2 | 18.26 | Unimer |
| 8 | TLL-poly(NIPAM) | 37 | 8.2 | 163.0 | 10139.864 +/- 4081.886 |
| 9 | TLL-poly(DPA) | 25 | 5.0 | 56.83 | 1034.588 +/- 385.152 |
| 10 | TLL-poly(DPA) | 25 | 8.2 | 142.4 | 11074.959 +/- 4122.937 |

Using our single particle microscopy we directly observed for the first time, to the best of our knowledge, the process of biohybrid assembly. We employed CALB-poly(NIPAM) which has a slower assembly rate compared to TLL-poly(NIPAM,) allowing us to observe real-time assembly process in multiple levels on TIRF experiments: firstly, we let the unimer or oligomeric CALB-poly(NIPAM) to be attached on a glass surface and incremented the temperature while recording with a frame rate of 2 frames per minute allowing for the direct observation of the assembly process (movie **S1**). Secondly, we incubated CALB-poly(NIPAM) at 37 °C in solution for 30 min to promote self-assembly before adding to the microscope slide and maintained constant heating at 37 °C for 1 hour. Under these experiment conditions, we were allowed to perform direct recording of the transition of lower order nanoparticles (heated at 37 °C for 30 min) to higher order nanoparticles (further heating for 1 hour) (Fig. **4B** and **C**). In this context, we assessed the number of enzymes in each assembled nanoparticle and its temporal variation. Figure **4D** displays four representative traces of time-dependent increase of the number of enzymes per nanoparticle, highlighting the previously unquantified heterogeneity of the process, since the final enzyme count per nanoparticle varies from 20k to 50k. Interestingly, a temporal heterogeneous growth that varies across traces is summarised in the mean increase that contains a large standard deviation across traces (Fig. **4E**).

**Figure 4:**
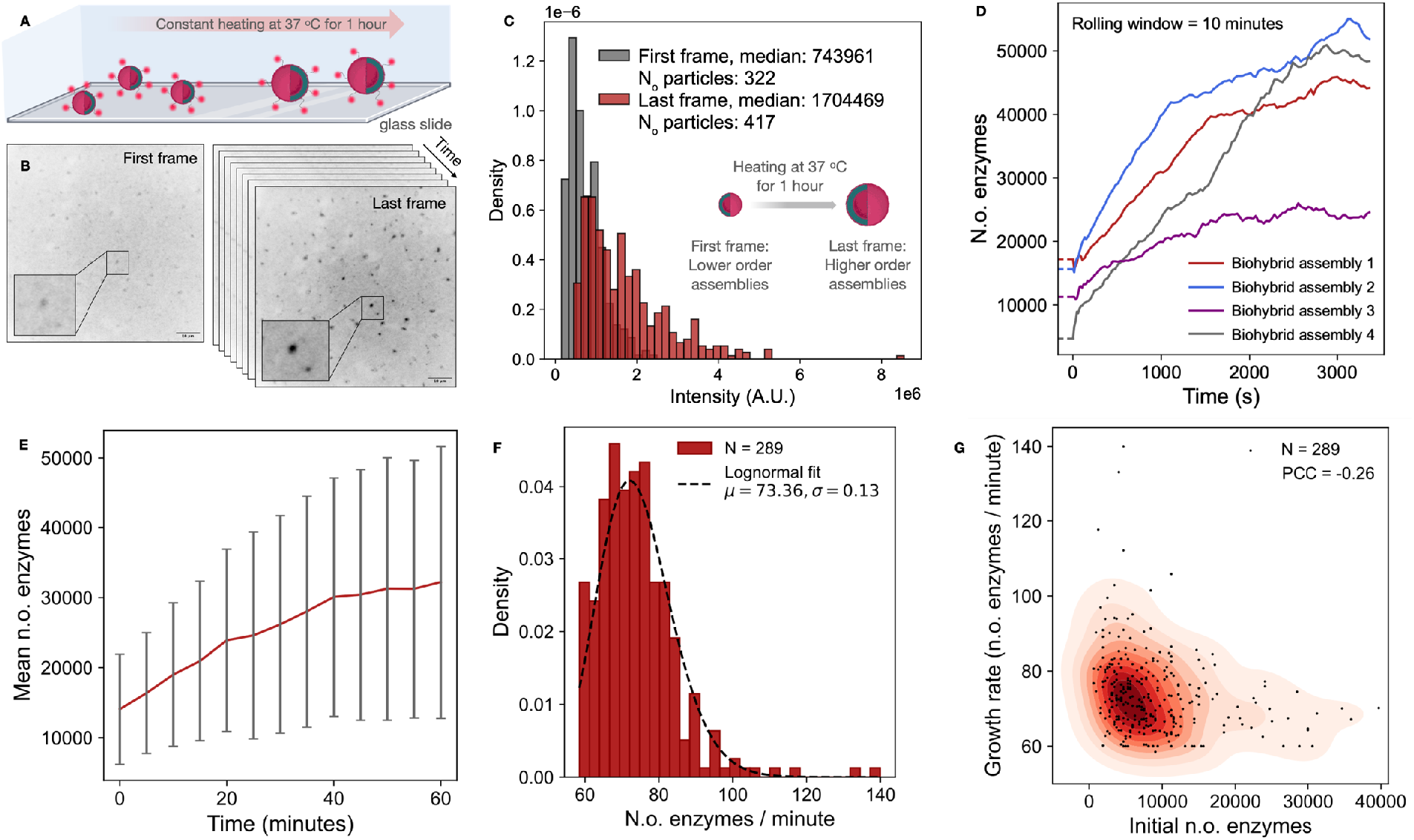
Real time direct recordings of the temperature dependent formation of higher order nanoparticles of CALB-poly(NIPAM) upon heating at 37 °C for 1h. **A**. Schematic representation of the TIRF microscope set up used for the direct recordings of higher order nanoparticle formation at the single particle level. **B**. Representative experimentally acquired TIRF images recorded upon heating the temperature responsive CALB-poly(NIPAM) showed the formation of higher order nanoparticles leading to intensity increase. **C**. Intensity distribution of the first and the last frame of a series of images recorded upon heating CALB-poly(NIPAM) at 37 °C for 1h. In the first frame the intensity distribution has median of 743960.6 with a sigma of 415778.9. The number of detected particles is 322. In the last frame the intensity distribution has median of 1704468.9 with a sigma of 1028895.9. The number of detected particles is 417. **D**. Four representative traces showing the time-depended increase of enzymes per nanoparticle upon heating. Note the stark heterogeneity in growth rates, initial and final enzyme numbers per nanoparticle that is revealed by the single particle readout. **E**. Quantification of the mean number of enzymes per CALB-poly(NIPAM) nanoparticle upon heating. Error bars correspond to the standard deviation of the mean of the number of enzymes for all the CALB-poly(NIPAM) particles every 5 min (10 frames). **F**. Growth rate distribution of the assembled CALB-poly(NIPAM). **G**. Scatter plot of the growth rate distribution vs initial number of the assembled CALB-poly(NIPAM). Pearson correlation coefficient (PCC) shows mild negative correlation.

We further examined whether the observed heterogeneity in growth rates stems from the existence of diverse biohybrid assembly dimensions. By tracking the assembly of each biohybrid over time, we were able to extract their individual growth rates, which showed to be lognormally distributed with a mean ∼73 enzymes per minute (Fig. **4F**). Figure **4G** reveals small or no correlation of the initial enzymes and their dimension to growth rate. This variation in maximum number and growth rate may originate from monomeric depletion or reaching maximum growth for each morphology type and may even depend on the spontaneous curvature of the biohybrid linking behaviour to protein-polymer conjugate spontaneous curvature^33^ (see supplementary movie).

### Single Particle Activity Studies of Enzyme Biohybrids

Multiple parameters including the synthetic approach and/or the assembly itself might induce loss of enzymatic activity of enzyme-polymer biohybrids challenging their implementation as tailored biocatalysts. We, therefore, quantified the catalytic activity of lipase-polymer conjugates in a range of temperatures (*see methods and supplementary information*). The pre-fluorescent substrate 5-(6)-carboxyfluorescein diacetate (CFDA) that upon hydrolysis yields the fluorescent product, carboxyfluorescein (CF^-^) was used as substrate.^39^ The high fluorescence yield of the products allows functional recording by UV, fluorescence in conventional ensemble measurements and offers reliable readout at the fundamental level of individual enzyme catalytic cycles.^39^ UV measurements revealed that most of the CALB and TLL based biohybrids retained significant fractions of the corresponding native lipase activities (Fig. **S29** and **S31**).

Interestingly, the temperature-responsive TLL-poly(NIPAM) biohybrid demonstrated a catalytic activity surpassing that of native TLL by a factor of 2.3 when measured at 25 °C (Fig. **5A**, **S32**). Upon heating TLL-poly(NIPAM) exhibited superior catalytic activity across all tested temperatures with the highest ratio of 2.8 observed at 37.5 and 40 °C (Fig. **5A**, **S32**). Our findings show that TLL-poly(NIPAM) biohybrid exhibits enhanced catalytic activity as compared to the native enzyme paving the way for the design of optimised biocatalysts.

**Figure 5.**
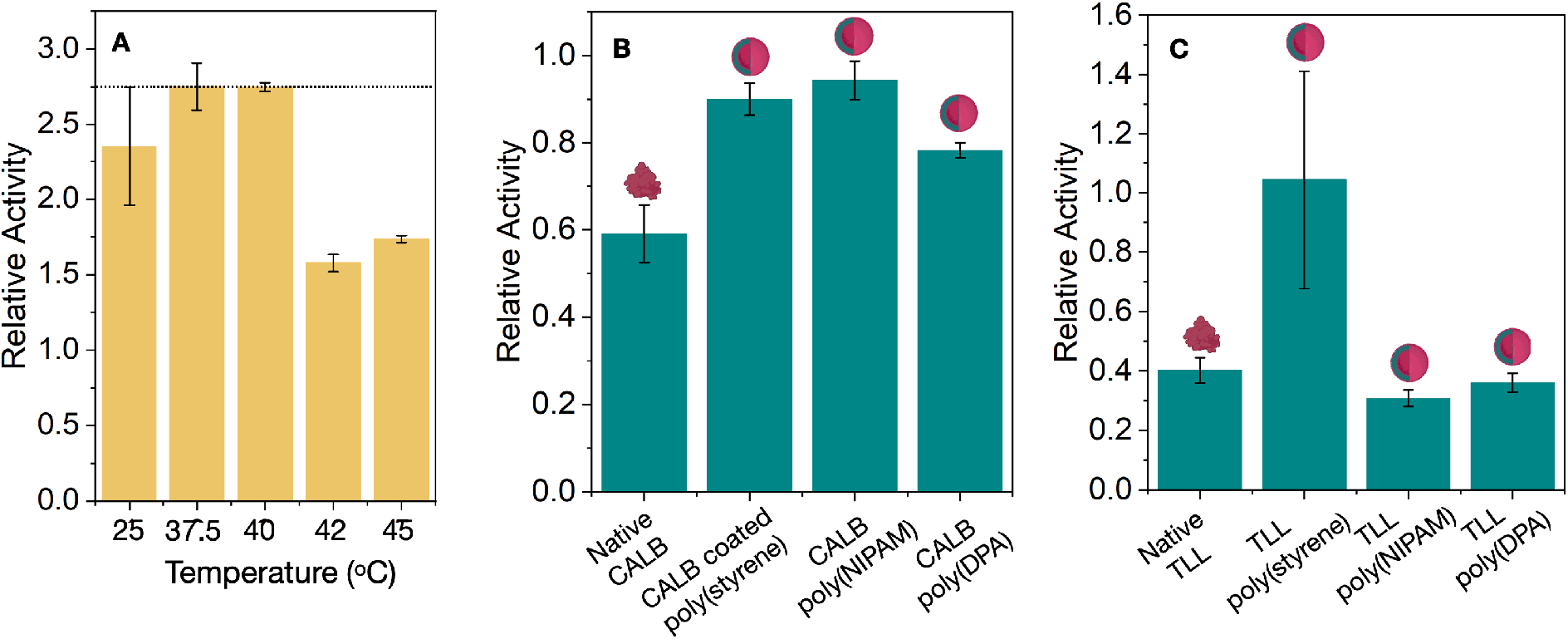
A. Relative activity of TLL-poly(NIPAM) as compared to native TLL at each tested temperature towards the pre-fluorescent CFDA substrate ester bonds hydrolysis to the fluorescent CF^-^ product. Error bars correspond to the standard deviation of 3 technical replicates of the catalytic activity measurement. **B**. Relative activity of 1 year old CALB samples stored at 4 °C as compared to fresh CALB samples (immediately after their preparation) on CFDA hydrolysis. **C**. Relative activity of 1 year old TLL samples stored at 4 °C as compared to fresh TLL samples (immediately after their preparation) on CFDA hydrolysis. Error bars correspond to the standard deviation of 3 technical replicates of the catalytic activity measurement.

A critical property of any biocatalytic system aimed for biotechnological applications is their long-term stability. To evaluate the long-term stability of CALB and TLL biohybrids we measured their activity on the CFDA substrate hydrolysis immediately after their preparation and after one year of storage. CALB and TLL biohybrids were found to lose less or similar fraction of their activity as compared to their native counterparts. Specifically, after one-year of storage or synthesis, native CALB retained only 60% of its initial catalytic efficiency while CALB-poly(NIPAM) retained 94% of its original activity, demonstrating a considerable improvement of stability (Fig. **5B**, **S37**). Following the same trend, the pH responsive CALB-poly(DPA) maintained 78% of its activity and CALB coated polystyrene reserved 90% of its activity (Fig. **5B**, **S37**). TLL biohybrids displayed similar stability, within error, to the native TLL enzyme upon storage for prolonged periods. However, in some cases, such as TLL-poly(styrene) higher stability was observed as compared to the native TLL (Fig. **5C**, **S38**).

To evaluate the potential correlation between enzymatic activity and biohybrid size, we developed a two-colour assay allowing the parallel recordings of the spatial localization dimensions and activity of hundreds of individual nanoparticles synchronously (*see methods and supplementary information*). In these simultaneous measurements, one channel outputs the biohybrid intensity and therefore size, while the second channel outputs the direct observation of product formation (Fig. **6A** and **B**). We, therefore, followed the catalytic activity of each biohybrid by measuring local intensities from the highly fluorescent product CF^-^, generated by enzymatic cleavage of the pre-fluorescent substrate CFDA. Simultaneously we measured the local background intensity, which enabled us to correct for systematic intensity changes not associated with enzymatic activity, such as laser power fluctuation, CFDA autohydrolysis, and spatial heterogeneities in the TIRF illumination profile (Fig. **S33** and **S34**).^39^ Each individual hydrolysis product (CF^-^) remains on the TLL or CALB active site for approximately 1 ms before diffusing away from the enzyme and escaping the TIRF detection volume.^39^ Although the photon output from a single catalytic event is typically insufficient to be detected using TIRF microscopy and exposure times of 50ms, the cumulative photon count from multiple of products produced from all the enzymes exceeds shot noise and can be detected. Average activities of the investigated CALB-poly(NIPAM) individual biohybrid followed a broad lognormal distribution (Fig **6C**). Control experiments in the absence of CFDA confirm the assay’s capacity to output the activity of individual biohybrids. Consistent with ensemble activity measurements, CALB biohybrids retained part of the CALB catalytic activity, among which the temperature responsive CALB-poly(NIPAM) at 37 °C, presented the highest catalytic activity. The same phenomenon was observed for TLL-poly(NIPAM) showing the highest catalytic activity amongst all TLL biohybrids (Fig. **S29**-**34**).

**Figure 6.**
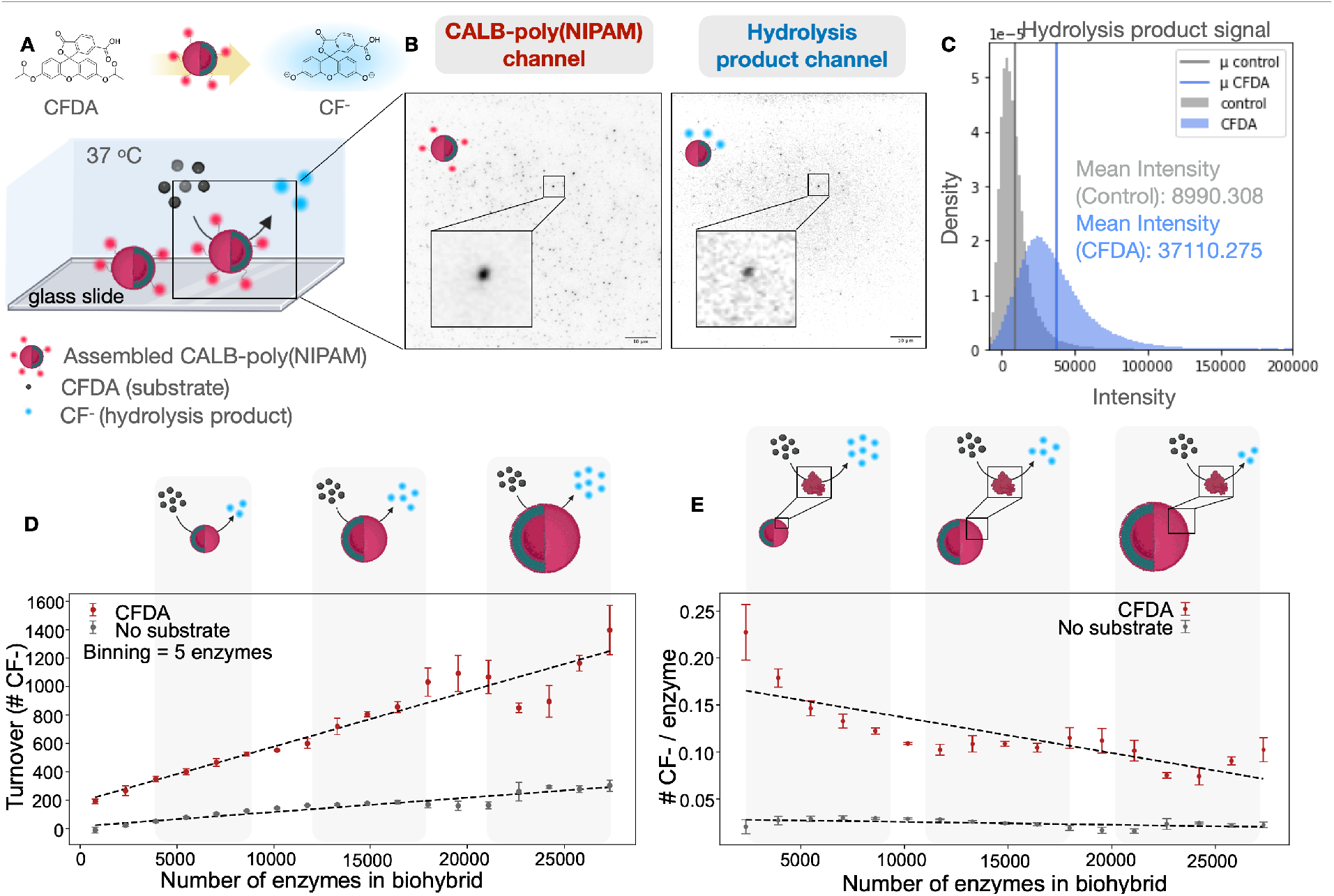
Quantification of activity of individual biohybrids and its dependence on nanoparticle dimensions **A**. Schematic representation of the direct recordings of catalytic activity of each individual CALB-poly(NIPAM) on CFDA substrate by TIRF microscopy. **B**. Representative experimentally TIRF images of the CALB-poly(NIPAM) (red) channel and the hydrolysis product (blue) channel. **C**. Distribution of background corrected intensities in the product (blue) channel in the presence and absence of (CFDA) (blue and grey graphs respectively). **D**. Quantification of number of turnovers per CALB-poly(NIPAM) nanoparticle at 37 °C in a population of nanoparticles of diverse dimensions (see SI with calibration curve). **E**. Calibrated number of turnovers per individual enzyme in each of the CALB-poly(NIPAM) nanoparticles at 37 °C in a population of nanoparticles with diverse dimensions.

Parallelised recordings of the dimensions and activity of hundreds of individual CALB-poly(NIPAM) nanoparticles synchronously allowed us to examine correlations between biohybrid size and enzymatic activity. Plotting the number of enzymatically produced CF^-^ products per nanoparticle showed as expected biohybrids of larger sizes, presented the larger number of turnovers (Fig. **6D**). This is to be expected as larger nanoparticles contain a higher number of enzymes. Interestingly plotting the number of turnovers per enzyme as part of the biohybrid revealed smaller biohybrids, although containing fewer enzymes, exhibit a small increase in turnover rates per enzyme. This may suggest that enzyme packing density in the biohybrid may influence the catalytic efficiency but delineating the precise origin of this activation is a subject of future investigations. Our findings not only provide insights into the fundamental aspect of enzyme behaviour in confined environments but also allow us to design enzyme-polymer nanoparticles of sizes that exhibit optimized operational efficiency of the constituent enzymes and improve the overall performance of the biohybrid detergents.

## CONCLUSIONS

A comprehensive investigation of the response, assembly, and activity of enzyme-polymer conjugates has revealed substantial enhancements in the stability and catalytic efficiency of CALB and TLL lipase based biohybrids, under a range of environmental conditions. A single particle assay allowed us to directly observe not only the multiple assembly pathways to form biohybrid spherical superstructures, but also the enzymatic turnovers of each assembly, providing new insights into constituent enzyme function efficiency. The quantification of the enzymatic turnovers revealed that smaller nanoparticles containing lower numbers of enzymes are constituted by enzymes of higher catalytic efficiency, highlighting their potential in sustainable industrial applications. The development of these biohybrids marks a remarkable advancement in the utilization of biocatalysts, offering both environmental and economic benefits. Future work will focus on optimizing these systems for broader industrial applications, aiming to fully harness the advantages of biohybrid technology. The development of these biohybrids marks a remarkable advancement in the utilization of biocatalysts, offering both environmental and economic benefits. Future work will focus on optimizing these systems for broader industrial applications, aiming to fully harness the advantages of biohybrid technology.

## EXPERIMENTAL METHODS

### CALB and TLL based biohybrids synthesis

The Eosin Y photocatalyzed oxygen tolerant grafting from polymerization approach was applied (*see Polymerization reaction in ESI*).^27^

### Biohybrid labelling

CALB and TLL biohybrids were labelled with ATTO 655 NHS-ester (ATTO-TEC) on free lysine residues by mixing a 5 mg/mL ATTO 655 NHS-ester solution in DMSO (3.9 μL, 22.5 × 10-6 mmol, 5.0 equiv.) with the 0.18 mM biohybrid solution in 20 mM phosphate buffer pH 8.2 (25 μM, 4.5 × 10^−6^, 1.0 equiv.) for 2 hours at room temperature. Free dyes were removed by dialysis using an 8-10 kDa MWCO membrane initially against a mixture of 1% DMSO in 5 mM phosphate buffer, then against 5 mM phosphate buffer, and finally against 20 mM phosphate buffer pH 8.2.

### Single particle assay for the assembling process

Ø25 mm round microscope glass slides were used for the surface preparation. Total Internal Reflection Fluorescence (TIRF) micro-scope (IX 83, Olympus) was used for the SM experiments. The laser line of 640 nm was used to excite the fluorophore ATTO-655 on the surface of the biohybrids. Imaging was performed using an exposure time of 50 ms, a penetration depth of 100 nm and an electron multiplying (EM) gain of 300. Each experimental run consisted of image series ranging from 60-120 frames using a laser line of 640 nm (2 mW) to excite ATTO-655 and a frame rate of 2 frames per minute.

### Single particle assay for enzymatic activity

Ø25 mm round microscope glass slides were used for the surface preparation. CFDA was used as the substrate. The hydrolysis product (CF^-^) is a fluorescent compound that has an emission peak at 517 nm. Total Internal Reflection Fluorescence (TIRF) microscope (IX 83, Olympus) was used for the SM experiments. The laser lines of 640 nm and 488 nm were used to excite the fluorophore ATTO-655 on the surface of the biohybrids and the hydrolysis product respectively. Biohybrids in 20 mM phosphate buffer pH 8.2 (10 nM in the final solution) were given 20 minutes to bind to the glass surface, followed by the addition of CFDA solution (30 nM in the final solution). Then, Imaging was performed with an exposure time of 50 ms, 100 nm penetration depth and 300 EM gain. Each image series contained 60 frames of the red channel (2 mW) and 60 frames of the blue channel (1.3 mW). The image series were recorded at 5-6 fields of view for each experiment with a frame rate 2 fpm. Image series under the same experimental condition were recorded in the absence of CFDA as a control experiment.

### Ensemble activity assay

Ensemble activity measurements were done on a Shimadzu UV-1900 UV-VIS spectrophotometer. 1 mg of 5-(6)-carboxyfluorescein diacetate (CFDA) was dissolved in 250 μL DMSO to form a stock solution which was fractionated and stored at -20 °C. 15 μL or 9 μL of a 0.31 mM CALB or 0,52 mM TLL solution respectively were diluted with 50 mM phosphate buffer pH 7.4 (2 mL) to form a 2.34 μM solution. The reaction was initiated by the addition of the 25 μL of the CFDA solution to the CALB solution and the esterase-like activity of CALB was monitored by UV at 453 nm at 37 °C. The ability of CALB- and TLL-biohybrids to hydrolyse CFDA was also tested following the same protocol. A blank experiment was performed using the same protocol in the absence of the enzyme. All reactions were performed in triplicates. The relative activities were calculated by dividing the slope of the CFDA hydrolysis by the biohybrid kinetics to the slope of the CFDA hydrolysis by native enzyme kinetics.

### Calculation of the number of enzymes

A single enzyme calibration curve was done by imaging native TLL labelled with ATTO-655. The laser line of 640 nm to excite the fluorophore (2 mW). Imaging was performed with an exposure time of 50, 100, 300, 400 and 500 nm, 100 nm penetration depth and 300 EM gain. The formula for linear regression was f(x)=237.54x + 4663.70. For 50 ms exposure time the native enzyme has a mean intensity f(50)=16340.70, indicating that CALB-poly(NIPAM) and TLL-poly(NIPAM) at 25 °C are at their monomeric form. Based on this we calculated the number of enzymes that the CALB and TLL biohybrids are composed of (*see single enzyme calibration curve in ESI*).

### Calculation of number of turnovers per enzyme

Glass slides 76 × 26 nm were dried with a nitrogen flow and activated using a plasma cleaner for 1 minute at a pressure between 300 mtorr to 400 mtorr. Immediately after activation the slides were attached to the glass slides to make flow cells. Each of the six champers was passivated with 80 μL of a PLL-g-PEG (1 mg/mL in HEPES buffer pH 5.5) and PLL-g-PEG-biotin (1 mg/mL in HEPES buffer pH 5.5) mixture in the ratio 100:1. The surface was incubated for 30 min. Excess of PLL-g-PEG mixture was removed by flushing five times with 50 μL HEPES buffer pH 5.5. 80 μL of the neutravidin solution (1 mg/mL in HEPES buffer pH 5.5) was added in each well. The surface was incubated for 15 min. Each well was washed with 20 mM PB buffer pH 8.2 at least 5 times before the sample addition. Biotinylated fluorescein (hydrolysis product) was tethered in the neutravidin surface and was imaged using the laser line of 488 nm to excite the hydrolysis product (1.3 mW). Imaging was performed with an exposure time of 50, 200, 400 and 500 nm, 100 nm penetration depth and 300 EM gain. The formula for linear regression was f(x)= 156.80x + 12670.83. Since CALB-poly(NIPAM) is a polydisperse sample, spherical structures of different sizes have a different catalytic activity outcome. According to our calibration curve, for 50 ms exposure time the hydrolysis product has a mean intensity f(50)=20510.83. Based on this we calculated the number of turnovers per biohybrid size and per enzyme in each biohybrid size for the CALB-poly(NIPAM) at 37 °C.

### Image Analysis

For the intensity extractions, an in-house algorithm based on Trackpy and Laplacian of Gaussian theory was used for analyzing .tif movies, detecting lipases and biohybrids on a surface and extracting intention values with in-house custom Python scripts for statical analysis of the results. The radius of the biohybrids in different scenarios was determined by formulas demonstrated in Quantifiable Information on a static image subsection. For each particle, the intensity within the ROI (defined by the σ of its detection) was integrated and adjusted for background noise.

## AUTHOR INFORMATION

## Author Contributions

N.S.H., K.V. and S.M.C. conceptualization and methodology; E.V., Th.L., J.B. and K.V. contributed to the investigation and data curation; E.V., E.W.S and A.O. data analysis; N.S.H., K.V. and S.M.C. supervision and resources; N.S.H. overall project management and supervision; E.V. and N.S.H. writing the original draft with support from E.W.S.; K.V. and S.M.C. review and editing; All authors have read and agreed to the published version of the manuscript.

## Notes

Sune M. Christensen is employed by Novonesis.

## ACKNOWLEDGMENT

We acknowledge Jeannette de Sparra Lundin, Gustav Hammerich Hansen and Christian Isak Jørgensen for assisting with CALB and TLL expression, purification, and characterization. The authors acknowledge funding from the Independent Research Fund Denmark (1127-00432B) Villum foundation (40801) and the NNF challenge center for Optimised Oligo Escape and control of disease (NNF23OC0081287).

