## Supplementary material for "Direct observation of assembly and function of trigger responsive lipase biohybrids"

a. Department of Chemistry, Nanoscience centre, University of Copenhagen, Denmark
b. Novo Nordisk center for Optimized Oligo Escape, University of Copenhagen, Denmark
c. Center for 4D cellular dynamics, Department of Chemistry, University of Copenhagen, Denmark
d. Department of Materials Science and Engineering, University of Crete, University Campus Voutes, Greece
e. Novonesis, Biologiens Vej 2, 2800 kgs, Lyngby, Denmark

| Table of Contents | Page |
|---|---|
| Materials | *S2* |
| Analytical Techniques | *S2* |
| Synthesis of the initiator 2-bromo-2-methyl-propionic acid 2-(2,5-dioxo-2,5-dihydro-pyrrol-1-yl)-ethyl ester (2) | *S4* |
| Synthesis of the Candida antarctica lipase B macroinitiator (CALB-Br) | *S5* |
| Synthesis of the Thermomyces lanuginosus lipase macroinitiator (TLL-Br) | *S6* |
| Polymerization Reactions | *S7* |
| CALB and TLL biohybrids labelling | *S12* |
| Characterization of CALB and TLL biohybrids | *S12* |
| Response and reversibility of the thermo-responsive CALB-poly(NIPAM) (bulk measurements) | *S19* |
| Response of the pH-responsive CALB-poly(DPA) (bulk measurements) | *S23* |
| Response and reversibility of the thermo-responsive TLL-poly(NIPAM) (bulk measurements) | *S24* |
| Response of the pH-responsive TLL-poly(DPA) (bulk measurements) | *S27* |
| Size distribution of CALB and TLL biohybrids | *S28* |
| Single enzyme calibration curve – $N_o$ enzymes in each biohybrid | *S33* |
| Response of CALB and TLL biohybrids at single particle level | *S35* |
| Catalytic activity of CALB and TLL biohybrids (bulk measurements) | *S37* |
| Catalytic activity of CALB and TLL biohybrids at single particle level | *S41* |
| Number of turnovers per enzyme in CALB-poly(NIPAM) at 37 $^oC$ | *S43* |
| Stability of CALB and TLL biohybrids | *S45* |
| CALB batch variation | *S46* |
| References | *S47* |

### 1. Materials

##### Materials

Chemicals such as sodium phosphate dibasic (CAS: 7558-79-4), sodium phosphate monobasic (CAS: 7778-77-0), sodium carbonate (CAS: 497-19-8), triethylamine (CAS: 121-44-8), *N*-hydroxysuccinimide (CAS: 6066-82-6), 2-bromo-2-methylpropionyl bromide (BIBB, CAS: 20769-85-1), 2',4',5',7'-tetrabromofluorescein (Eosin Y, CAS: 15086-94-9) and acetic acid were purchased from Sigma-Aldrich. Sodium acetate anhydrous was purchased from Merck. Monomers such as styrene (St, CAS: 100-42-5), *N*-isopropylacrylamide (NIPAM, CAS: 2210-25-5) and 2-(diisopropylamino)ethyl methacrylate (DPA, CAS: 16715-83-6) were purchased from Sigma-Aldrich. *N',N',N',N'*-tetramethylethylenediamine (TEMED, CAS: 110-18-9) was purchased from Bio-RAD. *Thermomyces lanuginosus Lipase* (TLL) and *Candida antarctica Lipase* B (CALB) WT enzymes were provided by Novonesis, Denmark. ATTO-655 NHS-ester was purchased from ATTO-TEC. CFDA used in bulk activity measurements was purchased from Invitrogen. CFDA used in microscopy studies was purchased from ThermoFisher. Biotin-4-fluorescein used for the calibration curve was purchased from Sigma Aldrich. Dialysis bags (Spectra/Por® Biotech Regenerated Cellulose Dialysis Membranes, MWCO 8-10 kDa) were purchased from Spectrum Labs. Dimethyl Sulfoxide (DMSO) was purchased from Merck, Darmstadt, Germany (CAS: 67-68-5).

##### Substrate surface preparation for single particle imaging

Ø25 mm round microscope glass slides were used.

### 2. Analytical Techniques

##### Irradiation Source

Blue LED flexible light strip, 60 LEDs/m, 10.8 w/m, 1000 lm/m.

##### Sodium dodecyl-sulfate polyacrylamide gel electrophoresis (SDS-PAGE)

PAGE electrophoresis was run using a 4% stacking gel and a 10% resolving gel. Samples mixed with an equal volume of electrophoresis sample buffer (125 mM Tris-HCl, pH 6.8, 5% SDS, 20% v/v glycerol, 0.004% bromophenol blue, 10% β-mercaptoethanol) and heated at 95 °C for 10 min prior to loading.

**UV-Vis Spectroscopy**

Activity and response studies were performed on a Shimadzu UV-1900 UV-VIS spectrophotometer.

**Id-3 spectraMax plate reader**

Activity measurements were also performed using an Id-3 spectraMax plate reader. Clear standard 96-well plates were used to measure absorbance at 453 nm.

**IR Spectroscopy**

Infrared spectroscopy was performed with a Nicolet 6700 Attenuated Total Reflection Fourier Transform Infrared (ATR FT-IR) spectrometer using Omnic (Thermo Electron Corporation) software.

**Scanning Electron Microscopy**

Scanning electron microscopy was performed with a JEOL JSM 6390LV Scanning Electron Microscope operated at 10-20 kV. SEM microscopy was also performed using a ZEISS Gemini SEM - Field Emission Scanning Electron Microscope. Before sample imaging, all samples were dried, and sputter coated with ca. 10 nm of gold (Au).

**NMR Spectroscopy**

$^{1}H$ and $^{13}C$ NMR spectra were recorded on Bruker 500 MHz spectrometer system. All chemical shifts are reported in ppm *(δ)* relative to tetramethylsilane, referenced to the chemical shifts of residual solvent resonances ($^{1}H$ and $^{13}C$). The following abbreviations were used to explain the multiplicities: s = singlet, bs = broad singlet, d = doublet, t = triplet, m = multiplet.

**DLS measurements**

The CALB and TLL biohybrids size distribution and response studies was determined by dynamic light scattering using a Malvern Zetasizer µV apparatus (Malvern Panalytical, UK) following the manufacturer's instructions. The sample was diluted to reach a final concentration of ~10 µM, filtered and its polydispersity index was measured at 25 or 37 °C. For the Refractive Index, protein was chosen. A QS High Precision Cell Cuvette made of Quartz SUPRASIL was used (Art. No. 105-231-001-8.5-40, light pate 1.25x1.25, center 8.5).

**Nanodrop**

A Thermo Scientific™ Invitrogen™ Nanodrop™ One Spectrophotometer with Qubit™ 4 Fluorometer was used for the labelling efficiency calculation of the labelled CALB and TLL biohybrods.

**Total internal reflection fluorescence (TIRF) microscopy**

A TIRF microscope (IX83, Olympus) was used for the single-particle tracking (SPT) experiments. Oil immersion objective (UAPON 100XOTIRF, NA 1.49, Olympus) and an EMCCD camera (ImagEM X2, Hamamatsu, Shizuoka, Japan) were used to record images and videos with a pixel width of 160 nm and field of view 81.92 µm × 81.92 µm. Laser lines of 640 nm and 488 nm were used to excite the fluorophores ATTO-655 (biohybrid surface) and $CF^-$ (activity measurements), respectively. Imaging was performed with an exposure time of 50 ms, 100 nm penetration depth, and 300 EM gain. Each image series contained 60-120 frames of the 488 nm channel (1.3 mW) and then 60-120 frames of the red channel (2 mW). Prior to imaging, 58 µL of 20 mM phosphate buffer pH 8.2 was added to the chamber. Then, 2 µL of the labeled biohybrids were added to the chamber, resulting in a final concentration of ~10 nM.

**Image Analysis**

For the intensity extractions, an in-house algorithm based on Trackpy and Laplacian of Gaussian theory was used for analyzing .tif movies, detecting lipases and biohybrids on a surface and extracting intention values with in-house custom Python scripts for statical analysis of the results.[1] The radius of the biohybrids in different scenarios was determined by formulas demonstrated in Quantifiable Information on a static image subsection.

#### 3. Experimental Procedures

##### 3.1 Synthesis of *N*-hydroxysuccinimide-2-bromo-2-methylpropionate (2)[2]

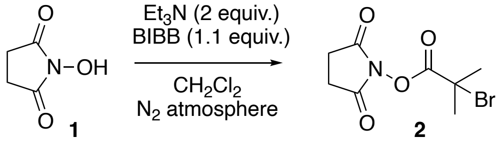


*N*-Hydroxysuccinimide (**1**, 575 mg, 5.0 mmol) and triethylamine ($Et_3N$, 1.4 mL, 10 mmol) were dissolved in 100 mL dichloromethane under nitrogen atmosphere, in a 250

ml round bottomed flask equipped with a magnetic stirrer. The flask was cooled to 0°C and 2-bromo-2-methylpropionyl bromide (BIBB, 0.68 mL, 5.5 mmol) was added dropwise. The mixture was stirred for 1 hour at 0 °C and allowed to reach room temperature for 2 hours. The reaction mixture was poured into an excess of cold water and extracted with diethyl ether (3 x 15 mL). The organic layer was washed with a saturated aqueous solution of sodium carbonate (3 x 15 mL), diluted HCl aqueous solution (pH 4.5, 3 x 10 mL), and again with saturated aqueous solution of sodium carbonate (3 x 15 mL). The organic layer was dried over anhydrous magnesium sulphate, filtered and the solvent removed under reduced pressure to afford *N*-hydroxysuccinimide-2-bromo-2-methylpropionate as a white solid (**2**, 498 g, 1.89 mmol, 38%).

**$^1$H NMR** (500 MHz, $CDCl_3$) $\delta$ = 2.87 (s, 4H, 2 x $COCH_2$), 2.08 (s, 6H, 2 x $CH_3$).

**$^{13}$C NMR** (125 MHz, $CDCl_3$) $\delta$ = 168.5, 167.5, 51.1, 30.7, 25.6.

##### 3.2 Synthesis of the Candida antarctica lipase B macroinitiator (CALB-Br)

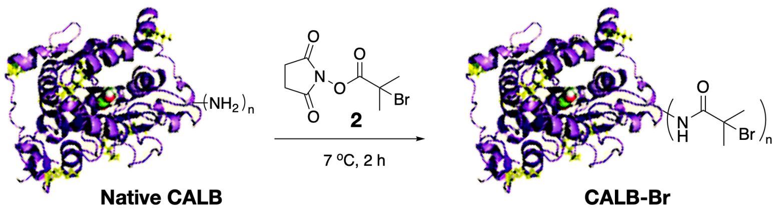


*N*-hydroxysuccinimide-2-bromo-2-methylpropionate (**2**) (3.2 mg, 0.012 mmol, 20 equiv.) in DMSO (40 µL) was slowly added to 2.0 mL of a 0.31 mM solution of native CALB (0.0006 mmol, 1.0 equiv.) in 100 mM phosphate buffer (pH 7.4). The reaction mixture was gently shaken for 2 hours at 7°C. The mixture was subsequently extensively dialyzed initially against 1% DMSO in 5 mM phosphate buffer pH 7.4 then against 5 mM phosphate buffer pH 7.4, and finally against 20 mM phosphate buffer pH 7.4 using a 10 kDa MWCO membrane. The resulting solution of CALB-macroinitiator (CALB-Br) was characterized with FT-IR and SDS-PAGE electrophoresis and stored at 4 °C until further use.

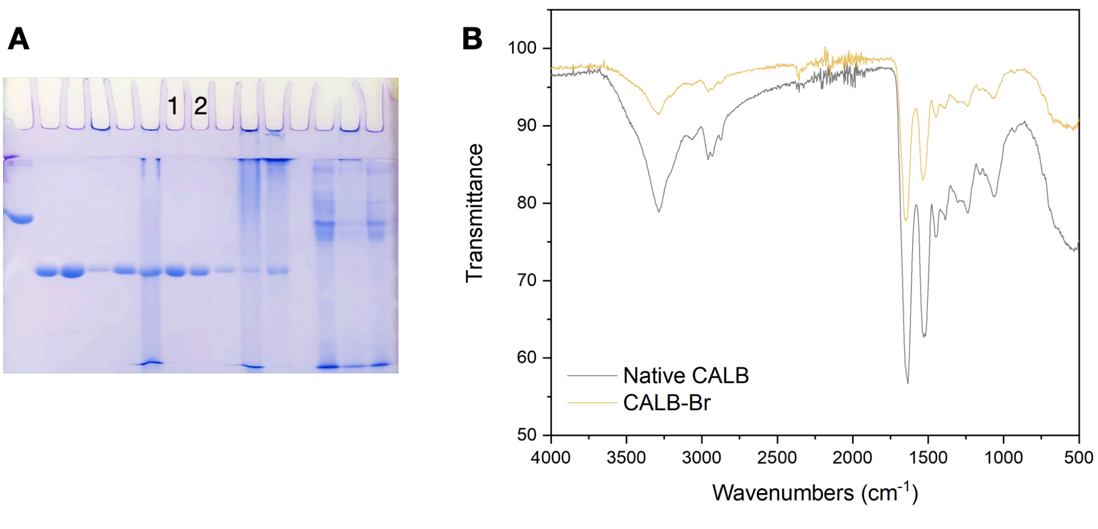


**Figure S1**. **A**. SDS-PAGE electrophoresis. Lane 1: native CALB, lane 2: CALB-Br. **B**. FT-IR spectra of CALB-Br and native CALB.

##### 3.3 Synthesis of the Thermomyces lanuginosus lipase macroinitiator (TLL-Br)

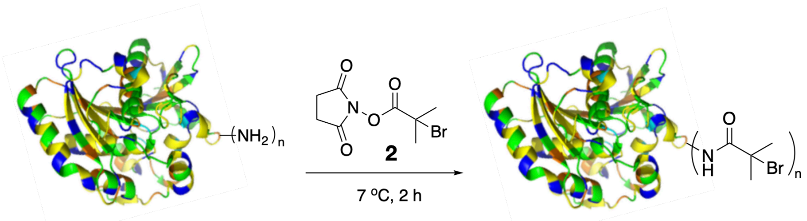


**Native TLL** **TLL-Br**

*N*-hydroxysuccinimide-2-bromo-2-methylpropionate (**2**) (5.5 mg, 0.02 mmol, 20 equiv.) in DMSO (40 µL) was slowly added to 2.0 mL of a 0.52 mM solution of native TLL (0.001 mmol, 1.0 equiv.) in 100 mM phosphate buffer (pH 7.4). The reaction mixture was gently shaken for 2 hours at 7°C. The mixture was subsequently extensively dialyzed initially against 1% DMSO in 5 mM phosphate buffer pH 7.4 then against 5 mM phosphate buffer pH 7.4, and finally against 20 mM phosphate buffer pH 7.4 using a 10 kDa MWCO membrane. The resulting solution of TLL-macroinitiator (TLL-Br) The resulting solution of CALB-macroinitiator (CALB-Br) was characterized with FT-IR SDS-PAGE electrophoresis and stored at 4 °C until further use.

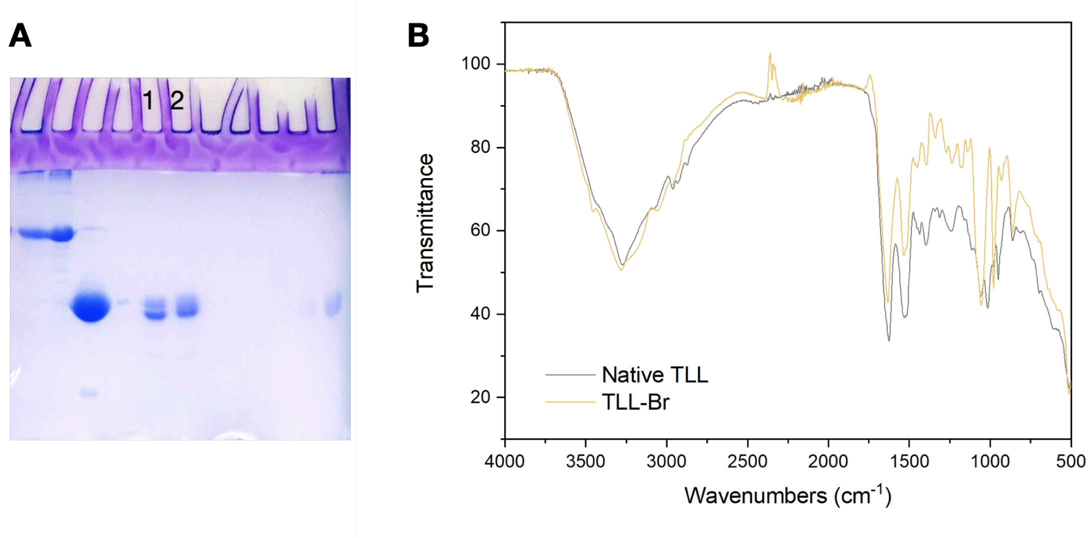


**Figure S2**. **A**. SDS-PAGE electrophoresis. Lane 1: native TLL, lane 2: TLL-Br. **B**. FT-IR spectra of TLL-Br and native TLL.

##### 3.4 Polymerization reactions

A stock solution of Eosin Y was prepared by dissolving Eosin Y (2.00 mg, 0.0293 mmol, 13 equiv.) in 20 mM phosphate buffer pH 7.4 (200 µL) with the aid of sonication for 3 minutes.

A stock solution of TEMED was prepared by dissolving TEMED (3.3 µL, 0.0218 mmol, 100 equiv.) in nanopure water (100 µL).

A ventilator was used to avoid temperature increase due to heating from blue LEDs, maintaining the temperature between 25 and 32 °C.

###### 3.4.1 Synthesis of lipase coated polymeric nanoparticles

**Experimental procedure of the Eosin Y catalyzed polymerization protocol of styrene for the synthesis of CALB coated poly(styrene) nanoparticles.**

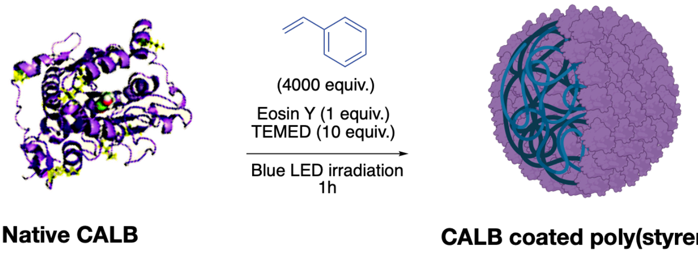


15 µL (0.218 µmol, 1.0 equiv.) of the Eosin Y stock solution and 10 µL (2.180 µmol, 10 equiv.) of the TEMED stock solution were dissolved in nanopure water to afford a solution with fixed total volume (450 µL). The emulsion of the monomer was formed by

adding the hydrophobic monomer styrene (100 µL, 874 µmol, 4000 equiv.) and sonicating for ca. 3 minutes. The resulting emulsion was immediately transferred to a 6 mL polypropylene syringe equipped with a stirring bar, containing a 0.31 mM solution of native CALB (0.696 mL, 0.218 µmol, 1.0 equiv.) in 20 mM phosphate buffer, pH 7.4. The headspace was eliminated to avoid the presence of undissolved oxygen and the reaction syringe was capped and placed under blue LED irradiation for 1 hour with moderate stirring. The reaction mixture was then dialyzed using a 10 kDa MWCO regenerated cellulose dialysis membrane initially against 5 mM phosphate buffer, pH 7.4, 1 % DMSO, then against 5 mM phosphate buffer, pH 7.4, and finally against 20 mM phosphate buffer, pH 7.4. The product solution was analyzed by means of native or SDS PAGE electrophoresis and FT-IR spectroscopy. Dilute suspensions of the product in nanopure water were imaged with SEM. The product was stored at 4 °C until further use.

##### 3.4.2 Grafting polymers from the surface of lipases

**Experimental procedure of the Eosin Y catalyzed polymerization protocol of NIPAM for the synthesis of CALB-poly(NIPAM) bioconjugates.**

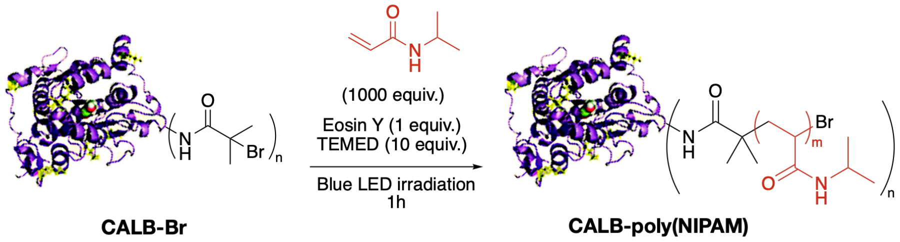


15 µL (0.218 µmol, 1.0 equiv.) of the Eosin Y stock solution and 10 µL (2.180 µmol, 10 equiv.) of the TEMED stock solution were dissolved in nanopure water to afford a solution with fixed total volume (450 µL). *N*-Isopropylacrylamide (NIPAM, 25 mg, 0.218 mmol, 1000 equiv.) was added in the Eosin Y and TEMED solution. The resulting mixture was immediately transferred to a 6 mL polypropylene syringe equipped with a stirring bar, containing a 0.31 mM solution of CALB-Br (0.696 mL, 0.218 µmol, 1.0 equiv.) in 20 mM phosphate buffer, pH 7.4. The headspace was eliminated to avoid the presence of undissolved oxygen and the reaction syringe was capped and placed under blue LED irradiation for 1 hour with moderate stirring. The reaction mixture was then dialyzed using a 10 kDa MWCO regenerated cellulose dialysis membrane initially against 5 mM phosphate buffer, pH 7.4, 1 % DMSO, then against 5 mM phosphate buffer, pH 7.4, and finally against 20 mM phosphate buffer, pH 7.4. The product

solution was analyzed by means of native or SDS PAGE electrophoresis and FT-IR spectroscopy. Dilute suspensions of the product in nanopure water were imaged with SEM and FE-SEM. The product was stored at 4 °C until further use.

**Experimental procedure of the Eosin Y catalyzed polymerization protocol of DPA for the synthesis of CALB-poly(DPA) bioconjugates.**

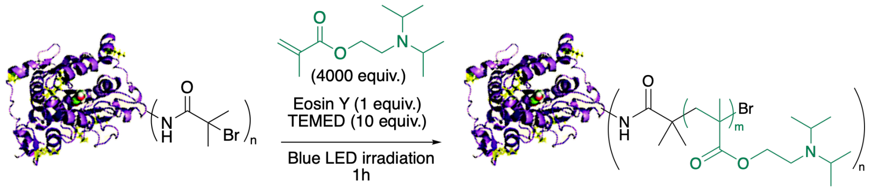


15 µL (0.218 µmol, 1.0 equiv.) of the Eosin Y stock solution and 10 µL (2.180 µmol, 10 equiv.) of the TEMED stock solution were dissolved in nanopure water to afford a solution with fixed total volume (450 µL). The emulsion of the monomer was formed by adding the hydrophobic monomer 2-(diisopropylamino)ethyl methacrylate (DPA, 207 µL, 0.874 mmol, 4000 equiv.) and sonicating for ca. 3 minutes. The resulting emulsion was immediately transferred to a 6 mL polypropylene syringe equipped with a stirring bar, containing a 0.31 mM solution of CALB-Br (0.696 mL, 0.218 µmol, 1.0 equiv.) in 20 mM phosphate buffer, pH 7.4. The headspace was eliminated to avoid the presence of undissolved oxygen and the reaction syringe was capped and placed under blue LED irradiation for 1 hour with moderate stirring. The reaction mixture was then dialyzed using a 10 kDa MWCO regenerated cellulose dialysis membrane initially against 5 mM phosphate buffer, pH 7.4, 1 % DMSO, then against 5 mM phosphate buffer, pH 7.4, and finally against 20 mM phosphate buffer, pH 7.4. The product solution was analyzed by means of native or SDS PAGE electrophoresis and FT-IR spectroscopy. Dilute suspensions of the product in nanopure water were imaged with SEM and FE-SEM. The product was stored at 4 °C until further use.

**Experimental procedure of the Eosin Y catalyzed polymerization protocol of styrene for the synthesis of TLL-poly(styrene) bioconjugates.**

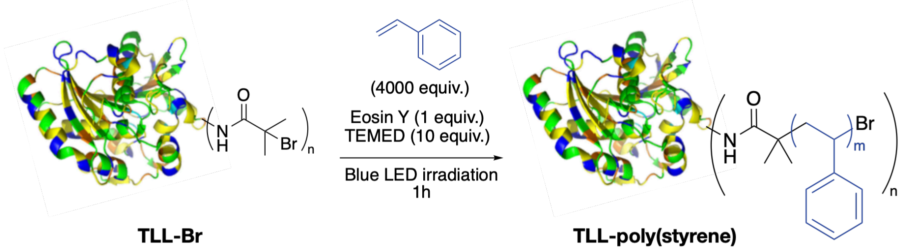


15 µL (0.218 µmol, 1.0 equiv.) of the Eosin Y stock solution and 10 µL (2.180 µmol, 10 equiv.) of the TEMED stock solution were dissolved in nanopure water to afford a solution with fixed total volume (450 µL). The emulsion of the monomer was formed by adding the hydrophobic monomer styrene (100 µL, 0.874 mmol, 4000 equiv.) and sonicating for ca. 3 minutes. The resulting emulsion was immediately transferred to a 6 mL polypropylene syringe equipped with a stirring bar, containing a 0.31 mM solution of TLL-Br (0.436 mL, 0.218 µmol, 1.0 equiv.) in 20 mM phosphate buffer, pH 7.4. The headspace was eliminated to avoid the presence of undissolved oxygen and the reaction syringe was capped and placed under blue LED irradiation for 1 hour with moderate stirring. The reaction mixture was then dialyzed using a 10 kDa MWCO regenerated cellulose dialysis membrane initially against 5 mM phosphate buffer, pH 7.4, 1 % DMSO, then against 5 mM phosphate buffer, pH 7.4, and finally against 20 mM phosphate buffer, pH 7.4. The product solution was analyzed by means of native or SDS PAGE electrophoresis and FT-IR spectroscopy. Dilute suspensions of the product in nanopure water were imaged with FE-SEM. The product was stored at 4 °C until further use.

**Experimental procedure of the Eosin Y catalyzed polymerization protocol of NIPAM for the synthesis of TLL-poly(NIPAM) bioconjugates.**

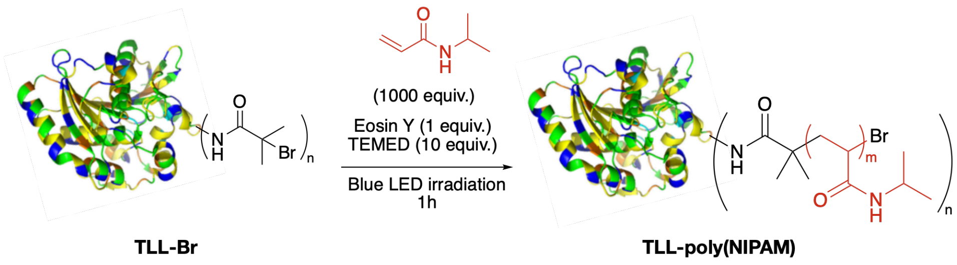


15 µL (0.218 µmol, 1.0 equiv.) of the Eosin Y stock solution and 10 µL (2.180 µmol, 10 equiv.) of the TEMED stock solution were dissolved in nanopure water to afford a

solution with fixed total volume (450 µL). *N*-Isopropylacrylamide (NIPAM, 25 mg, 0.218 mmol, 1000 equiv.) was added in the Eosin Y and TEMED solution. The resulting mixture was immediately transferred to a 6 mL polypropylene syringe equipped with a stirring bar, containing a 0.31 mM solution of TLL-Br (0.436 mL, 0.218 µmol, 1.0 equiv.) in 20 mM phosphate buffer, pH 7.4. The headspace was eliminated to avoid the presence of undissolved oxygen and the reaction syringe was capped and placed under blue LED irradiation for 1 hour with moderate stirring. The reaction mixture was then dialyzed using a 10 kDa MWCO regenerated cellulose dialysis membrane initially against 5 mM phosphate buffer, pH 7.4, 1 % DMSO, then against 5 mM phosphate buffer, pH 7.4, and finally against 20 mM phosphate buffer, pH 7.4. The product solution was analyzed by means of native or SDS PAGE electrophoresis and FT-IR spectroscopy. Dilute suspensions of the product in nanopure water were imaged with FE-SEM. The product was stored at 4 °C until further use.

**Experimental procedure of the Eosin Y catalyzed polymerization protocol of DPA for the synthesis of TLL-poly(DPA) bioconjugates.**

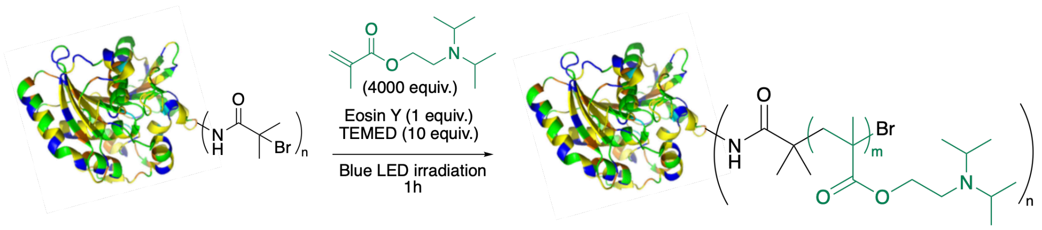


15 µL (0.218 µmol, 1.0 equiv.) of the Eosin Y stock solution and 10 µL (2.180 µmol, 10 equiv.) of the TEMED stock solution were dissolved in nanopure water to afford a solution with fixed total volume (450 µL). The emulsion of the monomer was formed by adding the hydrophobic monomer 2-(diisopropylamino)ethyl methacrylate (DPA, 207 µL, 0.874 mmol, 4000 equiv.) and sonicating for ca. 3 minutes. The resulting emulsion was immediately transferred to a 6 mL polypropylene syringe equipped with a stirring bar, containing a 0.31 mM solution of TLL-Br (0.436 mL, 0.218 µmol, 1.0 equiv.) in 20 mM phosphate buffer, pH 7.4. The headspace was eliminated to avoid the presence of undissolved oxygen and the reaction syringe was capped and placed under blue LED irradiation for 1 hour with moderate stirring. The reaction mixture was then dialyzed using a 10 kDa MWCO regenerated cellulose dialysis membrane initially against 5 mM phosphate buffer, pH 7.4, 1 % DMSO, then against 5 mM phosphate buffer, pH 7.4, and finally against 20 mM phosphate buffer, pH 7.4. The product

solution was analyzed by means of native or SDS PAGE electrophoresis and FT-IR spectroscopy. Dilute suspensions of the product in nanopure water were imaged with FE-SEM. The product was stored at 4 °C until further use.

##### 3.5 CALB and TLL biohybrids labelling

CALB and TLL biohybrids were labelled with ATTO 655 NHS-ester (ATTO-TEC) on free lysine residues by mixing a 5 mg/mL ATTO 655 NHS-ester solution in DMSO (3.9 µL, 22.5 x $10^{-6}$ mmol, 5.0 equiv.) with the 0.18 mM biohybrid solution in 20 mM phosphate buffer pH 8.2 (25 µM, 4.5 x $10^{-6}$, 1.0 equiv.) for 2 hours at room temperature. Free dyes were removed by dialysis using an 8-10 kDa MWCO membrane initially against a mixture of 1% DMSO in 5 mM phosphate buffer, then against 5 mM phosphate buffer, and finally against 20 mM phosphate buffer pH 8.2. Labelling efficiencies were characterized by Nanodrop. The labelled enzymes were stored in 10 µM aliquots of 15 µL at -20 °C. Each aliquot was only used once.

##### 3.6 Characterization of CALB and TLL biohybrids

###### 3.6.1 Characterization of CALB coated poly(styrene)

CALB coated poly(styrene) was characterized by means of SDS-PAGE electrophoresis that revealed a higher migration rate of CALB coated poly(styrene) than the native CALB (Fig. 3, **A**), IR spectroscopy that showed a characteristic peak of 698.7 $cm^{-1}$ that can be attributed to the C-H bending of the aromatic ring of poly(styrene) (Fig. 3, **B**), DLS that showed an average hydrodynamic diameter of 83.55 +/- 47.98 nm (PDI: 0.330) (Fig. 3, **C**), and SEM microscopy that revealed spherical structures with diameters ranging from 50 to 100 nm (Fig. 3, **D**).

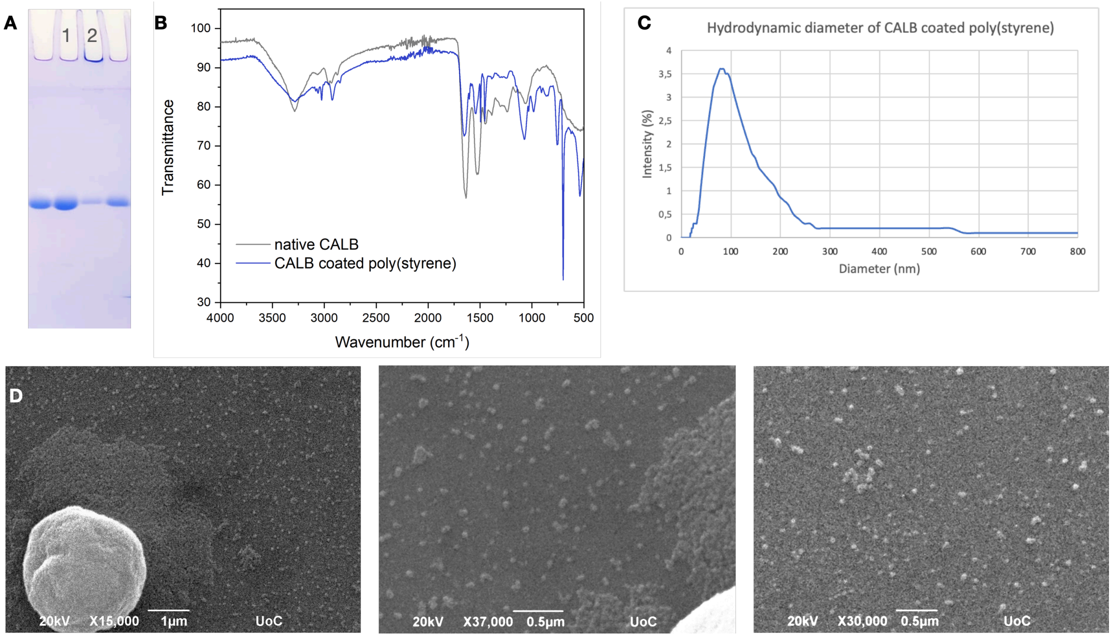


**Figure S3**. **A**. SDS-PAGE electrophoresis. Lane 1: native CALB, lane 2: CALB-poly(styrene). **B**. FT-IR spectra of CALB coated poly(styrene) and native CALB. **C**. Average hydrodynamic diameter distribution of CALB coated poly(styrene). **D**. SEM micrographs of CALB coated poly(styrene).

##### 3.6.2 Characterization of CALB-poly(NIPAM)

CALB-poly(NIPAM) was characterized by means of SDS-PAGE electrophoresis that revealed the consumption of CALB-Br (Fig. 4, **A** and **B**), $^{1}$H-NMR spectroscopy showed all the characteristic peaks of poly(NIPAM) (Fig. 4, **C**), SEM microscopy revealed spherical structures with diameters about 300 nm after 1hour rection time (Fig. 4, **D**), FE-SEM microscopy revealed spherical structures with diameters about 100 nm after 2 hours rection time (Fig. 4, **E**), DLS showed an average hydrodynamic diameter of 345.3 +/- 151.9 nm (PDI: 0.194) after 1 hour rection time (Fig. 1, **F**), IR spectroscopy that showed a peak at 1642.7 cm$^{-1}$ that can be attributed to the stretching of the C=O bond of amides, a peak at 1542.2 cm$^{-1}$ that can be attributed to the bending of N-H bond of amides, and two peaks at 1380.2 and 1363.4 cm$^{-1}$ that can be attributed to the stretching of C-N bond of poly(NIPAM) (Fig. 4, **G**).

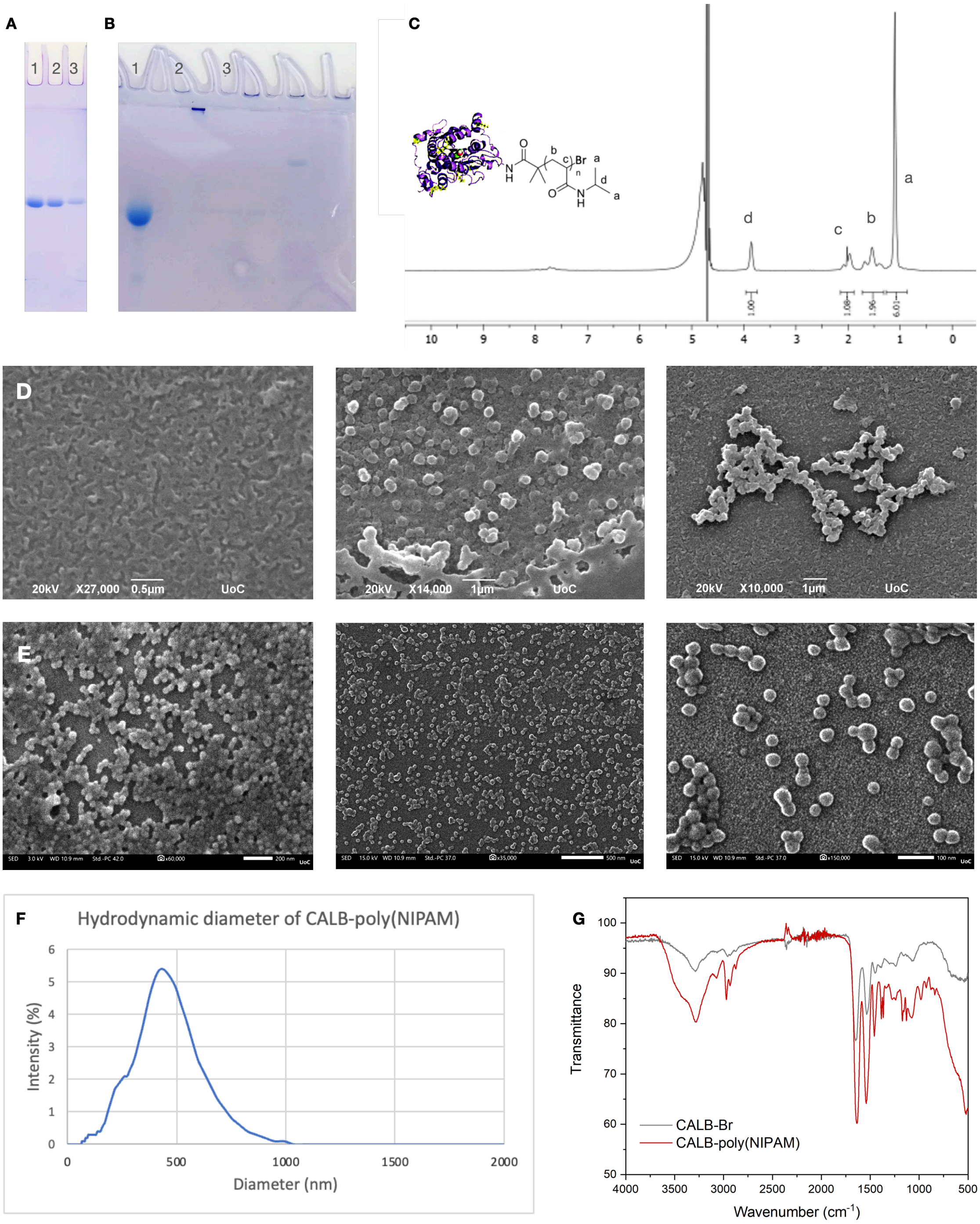


**Figure S4**. **A**. SDS-PAGE electrophoresis. Lane 1: native CALB, lane 2: CALB-Br, lane 3: CALB-poly(NIPAM) after 1 hour rection time. **B**. SDS-PAGE electrophoresis. Lane 1: native CALB, lane 2: CALB-poly(NIPAM) after 2 hours reaction time, lane 3: CALB-Br. **C**. $^1$H-NMR spectrum of CALB-poly(NIPAM) in $D_2O$ after water suppression. **D**. SEM micrographs of CALB-poly(NIPAM) after 1 hour reaction time. **E**. FE-SEM micrographs of CALB-poly(NIPAM) after 2 hours reaction time. **F**. Average hydrodynamic diameter distribution of CALB-poly(NIPAM) after 1 hour reaction time. **G**. FT-IR spectra of CALB-poly(NIPAM) and CALB-Br.

##### 3.6.3 Characterization of CALB-poly(DPA)

CALB-poly(DPA) was characterized by means of SDS-PAGE electrophoresis that revealed a higher migration rate of CALB-poly(DPA) than CALB-Br (Fig. 5, **A** and **B**), DLS showed an average hydrodynamic diameter of 87.71 +/- 25.30 nm (DPI: 0.083) (Fig. 5, **C**), IR spectroscopy showed a peak at 1730.5 cm$^{-1}$ that can be attributed to the stretching of the C=O bond of esters, and a peak at 1139.9 cm$^{-1}$ that can be attributed to the bending of N-H bond of tertiary amines of poly(DPA) (Fig. 5, **D**). SEM microscopy revealed spherical structures with diameters ranging from 50 to 100 nm after 1 hour reaction time (Fig. 5, **E**), FE-SEM microscopy revealed spherical structures with diameters ranging from 50 to 100 nm after 2 hours reaction time (Fig. 5, **F**).

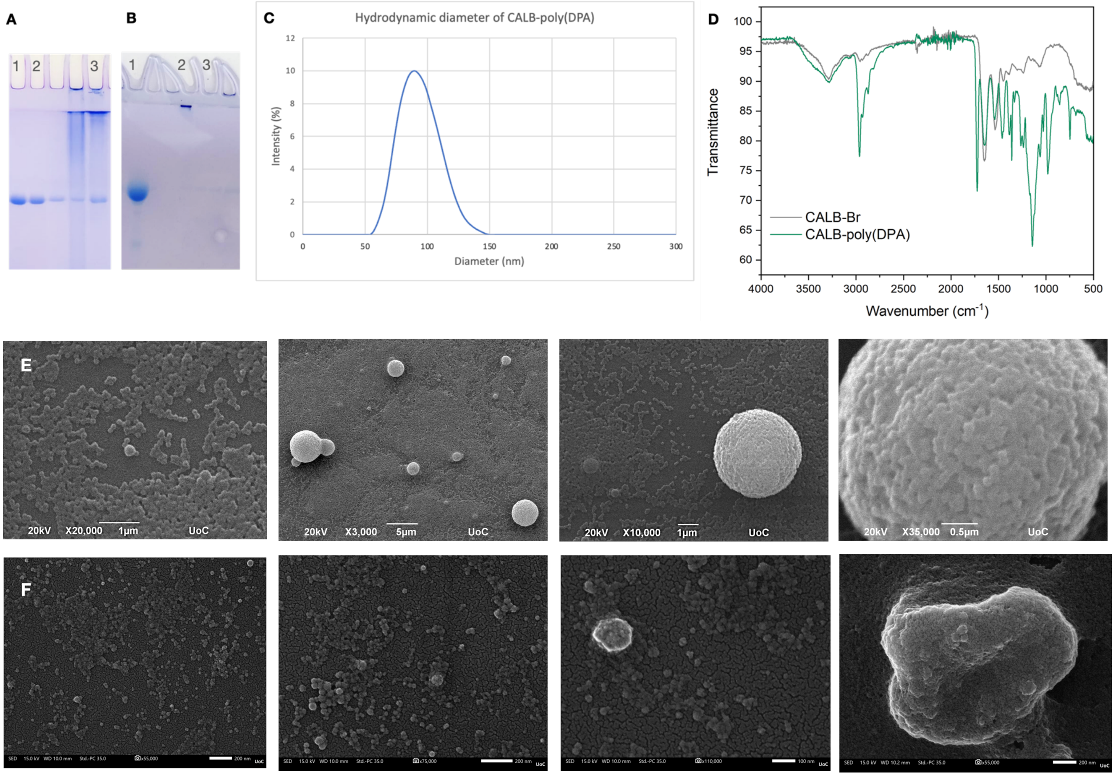


**Figure S5**. **A**. SDS-PAGE electrophoresis. Lane 1: native CALB, lane 2: CALB-Br, lane 3: CALB-poly(DPA) after 1 hour reaction time. **B**. SDS-PAGE electrophoresis. Lane 1: native CALB, lane 2: CALB-poly(DPA) after 2 hours reaction time, lane 3: CALB-Br. **C**. Average hydrodynamic diameter distribution of CALB-poly(DPA). **D**. FT-IR spectra of CALB-poly(DPA) and CALB-Br. **E**. SEM micrographs of CALB-poly(DPA) after 1 hour reaction time. **F**. FE-SEM micrographs of CALB-poly(DPA) after 2 hours reaction time.

##### 3.6.4 Characterization of TLL-poly(styrene)

TLL-poly(styrene) was characterized by means of SDS-PAGE electrophoresis that revealed a higher migration rate of TLL-poly(styrene) than the TLL-Br (Fig. 6, **A** and **B**), DLS that showed an average hydrodynamic diameter of 89.27 +/- 39.19 nm (PDI: 0.193) (Fig. 6, **C**), IR spectroscopy that showed a characteristic peak of 693.1 cm$^{-1}$ that can be attributed to the C-H bending of the aromatic ring of poly(styrene) (Fig. 6, **D**), FE-SEM microscopy revealed spherical structures with diameters ranging from 50 to 100 nm after 1 hour reaction time and using 4000 equiv. styrene (Fig. 6, **E**), FE-SEM microscopy revealed spherical structures with diameters ranging from 100 to 200 nm after 2 hours reaction time and using 6874 equiv. of styrene (Fig. 6, **F**).

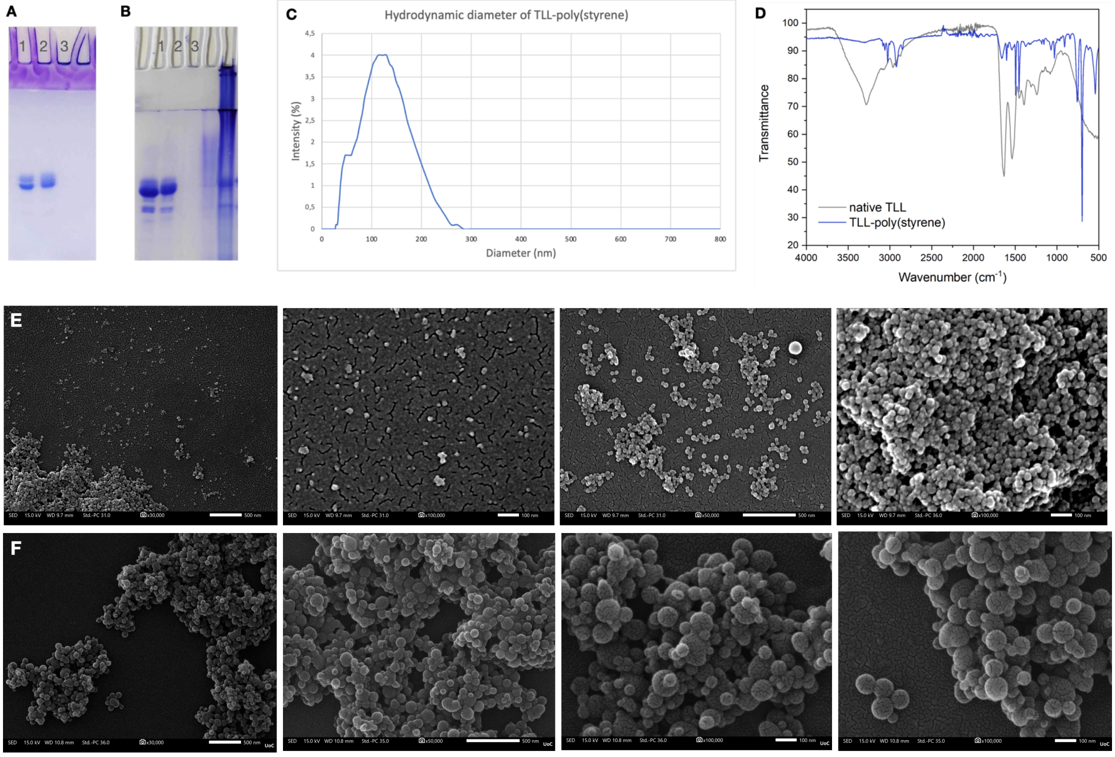


**Figure S6**. **A**. SDS-PAGE electrophoresis. Lane 1: native TLL, lane 2: TLL-Br, lane 3: TLL-poly(styrene) after 1 hour reaction time and using 4000 equiv. styrene. **B**. SDS-PAGE electrophoresis. Lane 1: native TLL, lane 2: TLL-Br, lane 3: TLL-poly(styrene) after 2 hours reaction time and using 6874 equiv. styrene. **C**. Average hydrodynamic diameter distribution of TLL-poly(styrene) after 1 hour reaction time and using 4000 equiv. styrene. **D**. FT-IR spectra of TLL-poly(styrene) and native TLL. **E**. FE-SEM micrographs of TLL-poly(styrene) after 1 hour reaction time and using 4000 equiv. styrene. **F**. FE-SEM micrographs of TLL-poly(styrene) after 2 hours reaction time and using 6874 equiv. styrene.

##### 3.6.5 Characterization of TLL-poly(NIPAM)

TLL-poly(NIPAM) was characterized by means of SDS-PAGE electrophoresis that revealed the consumption of TLL-Br (Fig. 7, **A** and **B**), $^{1}$H-NMR spectroscopy showed all the characteristic peaks of poly(NIPAM) (Fig. 7, **C**).IR spectroscopy that showed a peak at 1650.7 cm$^{-1}$ that can be attributed to the stretching of the C=O bond of amides, a peak at 1536.6 cm$^{-1}$ that can be attributed to the bending of N-H bond of amides, and two peaks at 1378.6 and 1359.4 cm$^{-1}$ that can be attributed to the stretching of C-N bond of poly(NIPAM) (Fig. 7, **D**). DLS showed an average hydrodynamic diameter of 163.0 +/- 27.13 nm (PDI: 0.028) after 1 hour reaction time and using 1000 equiv. NIPAM (Fig. 7, **E**). FE-SEM microscopy revealed spherical structures with diameters about 150 nm after 1 hour reaction time and using 1000 equiv. NIPAM (Fig. 7, **F**). FE-SEM microscopy revealed spherical assemblies with diameters about 0.5 to 1 µm after 2 hours reaction time and using 1740 equiv. NIPAM (Fig. 7, **G**).

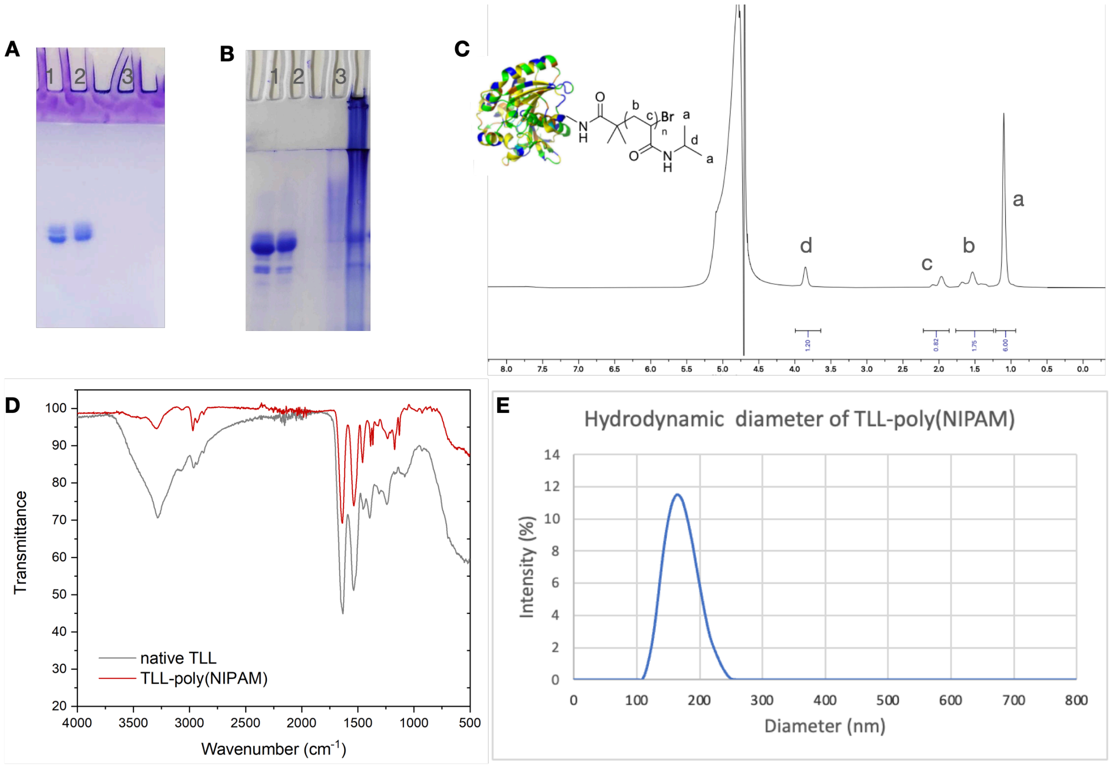

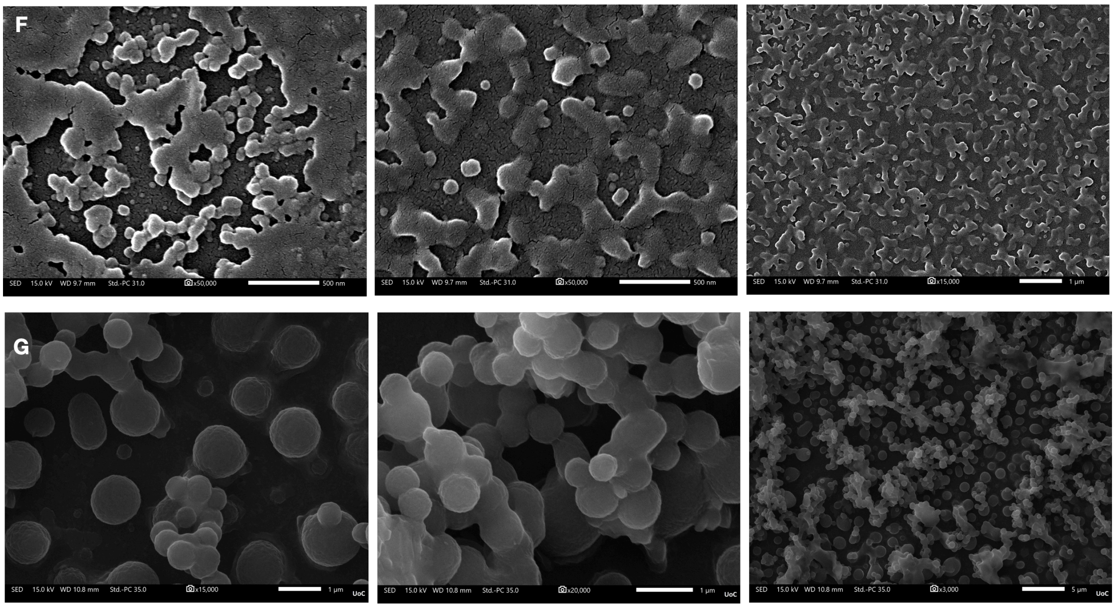


**Figure S7**. **A**. SDS-PAGE electrophoresis. Lane 1: native TLL, lane 2: TLL-Br, lane 3: TLL-poly(NIPAM) after 1 hour reaction time and using 1000 equiv. NIPAM. **B**. SDS-PAGE electrophoresis. Lane 1: native TLL, lane 2: TLL-Br, lane 3: TLL-poly(NIPAM) after 2 hours reaction time and using 1740 equiv. NIPAM. **C**. $^1$H-NMR spectrum of TLL-poly(NIPAM) in $D_2O$ after water suppression. **D**. FT-IR spectra of TLL-poly(NIPAM) and native TLL. **E**. Average hydrodynamic diameter distribution of TLL-poly(NIPAM) after 1 hour reaction time and using 1000 equiv. NIPAM. **F**. FE-SEM micrographs of TLL-poly(NIPAM) after 1 hour reaction time and using 1000 equiv. NIPAM. **G**. FE-SEM micrographs of TLL-poly(NIPAM) after 2 hours reaction time and using 1740 equiv. NIPAM.

##### 3.6.6 Characterization of TLL-poly(DPA)

TLL-poly(DPA) was characterized by means of SDS-PAGE electrophoresis that revealed a higher migration rate of TLL-poly(DPA) than TLL-Br (Fig. 8, **A** and **B**), IR spectroscopy showed a peak at 1731.3 $cm^{-1}$ that can be attributed to the stretching of the C=O bond of esters, and a peak at 1145.6 $cm^{-1}$ that can be attributed to the bending of N-H bond of tertiary amines of poly(DPA) (Fig. 8, **C**). DLS showed an average hydrodynamic diameter of 142.4 +/- 38.99 nm (DPI: 0.075) (Fig. 6, **D**), FE-SEM microscopy revealed spherical structures with diameters ranging from 50 to 100 nm (Fig. 8, **E**) after 1 hour reaction time and using 4000 equiv. DPA, and FE-SEM microscopy revealed spherical structures with diameters ranging from 50 to 100 nm (Fig. 8, **F**) after 2 hours reaction time and using 6874 equiv. DPA.

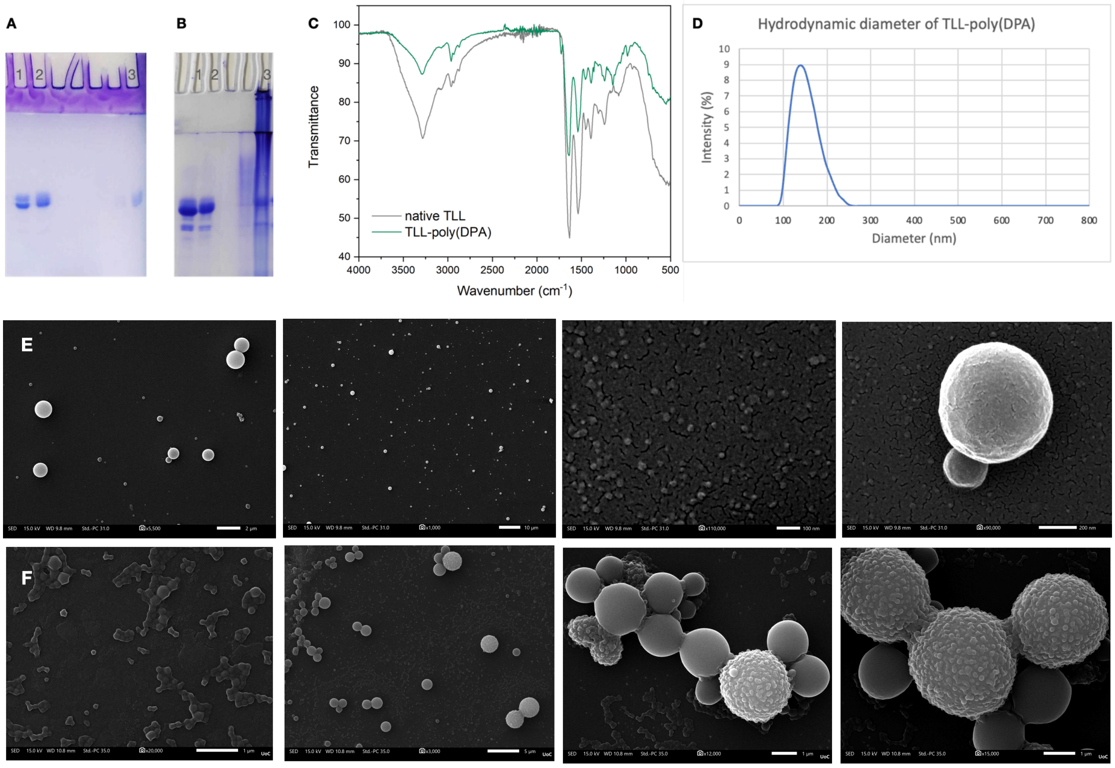


**Figure S8**. **A**. SDS-PAGE electrophoresis. Lane 1: native TLL, lane 2: TLL-Br, lane 3: TLL-poly(DPA) after 1 hour reaction time and using 4000 equiv. DPA. **B**. SDS-PAGE electrophoresis. Lane 1: native TLL, lane 2: TLL-Br, lane 3: TLL-poly(DPA) after 2 hours reaction time and using 6874 equiv. DPA. **C**. FT-IR spectra of TLL-poly(DPA) and native TLL **D**. Average hydrodynamic diameter distribution of TLL-poly(DPA) after 1 hour reaction time and using 4000 equiv. DPA. **E** FE-SEM micrographs of TLL-poly(DPA) ) after 1 hour reaction time and using 4000 equiv. DPA. **F** FE-SEM micrographs of TLL-poly(DPA) ) after 2 hours reaction time and using 6874 equiv. DPA.

#### 3.7 Response of CALB and TLL biohybrids (bulk measurements)

##### 3.7.1 Response and reversibility of the thermo-responsive CALB-poly(NIPAM)

The response and the reversibility of CALB-poly(NIPAM) was tested on UV-VIS spectrophotometer. Absorbance of CALB-poly(NIPAM) (9 μM) in 20 mM phosphate buffer solution (pH=7.4) was measured at 600 nm. CALB-poly(NIPAM) presented transmittance when was heated from 25-35 $^{o}$C. The transmittance was decreased with further heating up to 37.5, 40 and 44.5 $^{o}$C. The experiment was repeated successfully for multiple cycles, indicating that CALB-poly(NIPAM) maintains its responsiveness.

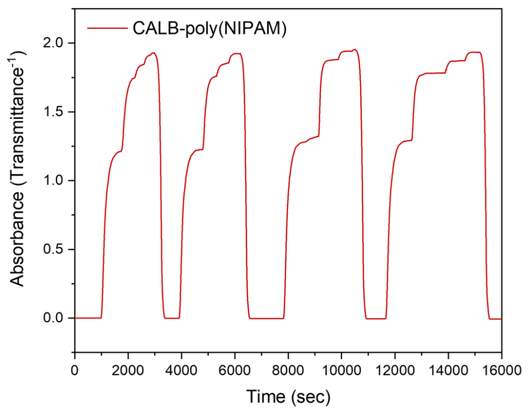


**Figure S9.** Transmittance of CALB-poly(NIPAM) in 20 mM phosphate buffer.

The responsiveness and the reversibility of CALB-poly(NIPAM) was also tested on DLS. The average hydrodynamic diameter (nm) of CALB-poly(NIPAM) was measured at 25 °C and after heating the sample at 37 °C back-and-forth five times. The average hydrodynamic diameter of CALB-poly(NIPAM) was measured from 12.31 to 19.54 nm at 25 °C and from 274.7 to 327.4 nm at 37 °C in all cases, indicating that CALB-poly(NIPAM) maintains its responsiveness.

**Table S1.** Average hydrodynamic diameter of CALB-poly(NIPAM) at 25 and 37 °C

| Cycle | Temperature (°C) | Size (d. nm) | PDI (%) |
|---|---|---|---|
| 1 | 25 | 13.50 ± 1.557 | 11.2 |
| | 37 | 327.4 ± 61.44 | 19.2 |
| 2 | 25 | 19.54 ± 5.956 | 28.9 |
| | 37 | 312.6 ± 66.75 | 21.1 |
| 3 | 25 | 12.31 ± 2.671 | 20.7 |
| | 37 | 289.0 ± 60.60 | 20.8 |
| 4 | 25 | 13.50 ± 3.158 | 21.9 |
| | 37 | 298.5 ± 71.99 | 22.8 |
| 5 | 25 | 12.89 ± 3.225 | 23.2 |
| | 37 | 274.7 ± 62.29 | 21.9 |

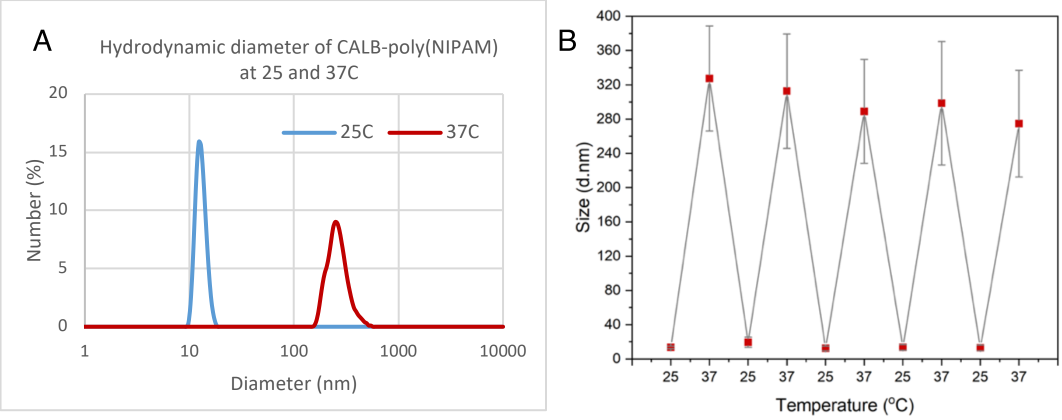


**Figure S10.** Reversibility of CALB-poly(NIPAM) in 20 mM phosphate buffer on DLS. **A**. The hydrodynamic diameter distribution of CALB-poly(NIPAM) at 25 and 37 °C. **B**. The average hydrodynamic diameter of CALB-poly(NIPAM) was measured at 25 °C and after heating the sample at 37 °C back-and-forth five times.

The average hydrodynamic diameter (nm) of CALB-poly(NIPAM) was also measured at different temperatures (20, 25, 30, 35, 37, 40 42 and 45 °C). The average hydrodynamic diameter detected on DLS was constantly increased by temperature, indicating that an assembly process is taking place at higher temperatures.

**Table S2.** Average hydrodynamic diameter of CALB-poly(NIPAM) heated at different temperatures

| Entry | Temperature (°C) | Size (d. Nm) | PDI (%) |
|---|---|---|---|
| **1** | 20 | 8.909 ± 1.667 | 16.1 |
| **2** | 25 | 7.406 ± 1.937 | 19.6 |
| **3** | 30 | 16.24 ± 6.430 | 28.9 |
| **4** | 35 | 236.9 ± 54.30 | 19.7 |
| **5** | 37 | 342.9± 64.59 | 17.6 |
| **6** | 40 | 359.1 ± 40.32 | 10.6 |
| **7** | 42 | 473.0 ± 103.1 | 19.5 |
| **8** | 45 | 787.6 ± 99.75 | 12.5 |

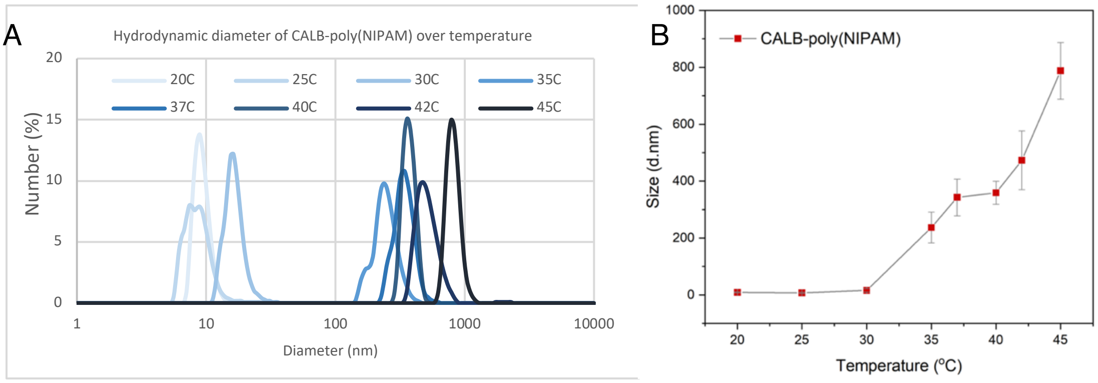


**Figure S11.** Response of CALB-poly(NIPAM) in 20 mM phosphate buffer on DLS. **A.** The hydrodynamic diameter distribution of CALB-poly(NIPAM) at 20, 25, 30, 35, 37, 40, 42 and 45 °C. **B**. The increase of the average hydrodynamic diameter of CALB-poly(NIPAM) over temperature.

The average hydrodynamic diameter (nm) of CALB-poly(NIPAM) was measured over time under constant heating at 37 °C. The size of CALB-poly(NIPAM) was increased I the first 5 min from 226.2 nm up to 412.5 nm and remained stable with further heating the sample at 37 °C.

**Table S3.** Average hydrodynamic diameter of CALB-poly(NIPAM) heated at 37 °C over time

| Entry | Incubation time (min) | Size (d. nm) | PDI (%) |
|---|---|---|---|
| 1 | 5 | 226.2 ± 108.1 | 21.0 |
| 2 | 10 | 412.5 ± 153.6 | 21.1 |
| 3 | 15 | 432.0 ± 170.2 | 18.9 |
| 4 | 20 | 452.4 ± 170.0 | 16.5 |

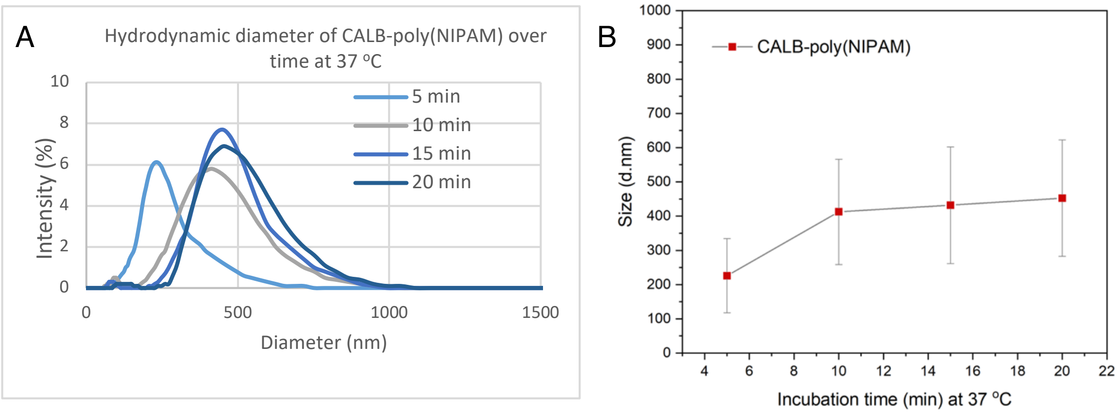


**Figure S12.** Size increase of CALB-poly(NIPAM) with constant heating at 37 °C over time measured on DLS. **A.** The hydrodynamic diameter distribution of CALB-poly(NIPAM) after 5, 10, 15 and 20 min of incubation at 37 °C. **B**. The increase of the average hydrodynamic diameter of CALB-poly(NIPAM) at each time point.

##### 3.7.2 Response of the pH-responsive CALB-poly(DPA)

The response of CALB-poly(DPA) was tested on UV-VIS spectrophotometer. The absorption spectra of CALB-poly(DPA) (9 µM) in 20 mM phosphate buffer solution at several pH values were recorded. The absorbance of CALB-poly(DPA) was detected at 600 nm. CALB-poly(DPA) presented low transmittance at higher pH values. The transmittance was increased by lowering the pH up to 5.9 adding aqueous HCl 1M and decreased by increasing the pH adding aqueous NaOH 1M. The experiment was repeated for 3 cycles.

**Table S4.** Absorbance at 600 nm of CALB-poly(NIPAM) at different pH values

| Entry | pH value | C (µM) | Abs at 600 nm | Corrected Abs at 9 µM |
|---|---|---|---|---|
| 1 | 7.48 | 9.00 | 0.61061 | 0.61061 |
| 2 | 7.20 | 8.95 | 0.53252 | 0.53549 |
| 3 | 6.98 | 8.91 | 0.49275 | 0.49773 |
| 4 | 6.70 | 8.86 | 0.38278 | 0.38883 |
| 5 | 6.42 | 8.82 | 0.16922 | 0.17267 |
| 6 | 5.97 | 8.78 | 0 | 0 |
| 7 | 6.46 | 8.69 | 0.01304 | 0.01351 |
| 8 | 6.79 | 8.61 | 0.01593 | 0.01665 |
| 9 | 7.13 | 8.53 | 0.04362 | 0.04602 |
| 10 | 5.60 | 8.41 | 0.00024 | 0.00025 |
| 11 | 6.65 | 8.29 | 0.01153 | 0.01252 |
| 12 | 7.42 | 8.18 | 0.02730 | 0.03003 |

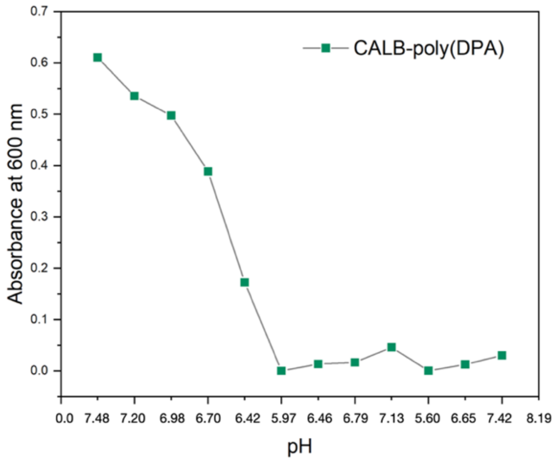


**Figure S13.** Transmittance of CALB-poly(DPA) in 20 mM phosphate buffer at different pH values.

The response of CALB-poly(DPA) was also tested on DLS. The average hydrodynamic diameter (nm) of CALB-poly(DPA) was also measured at different pH values (4, 5, 6, 7 and 8). The average hydrodynamic diameter detected on DLS was constantly increased by pH, indicating that an assembly process is taking place at higher pH values.

**Table S5.** Average hydrodynamic diameter of CALB-poly(DPA) at different pH values

| Entry | pH value | Size (d. nm) | PDI (%) |
|---|---|---|---|
| **1** | 4 | 8.507 ± 1.194 | 13.8 |
| **2** | 5 | 34.02 ± 4.523 | 35.8 |
| **3** | 6 | 129.9 ± 32.42 | 24.5 |
| **4** | 7 | 206.2 ± 55.05 | 25.1 |
| **5** | 8 | 236.9 ± 64.93 | 26.3 |

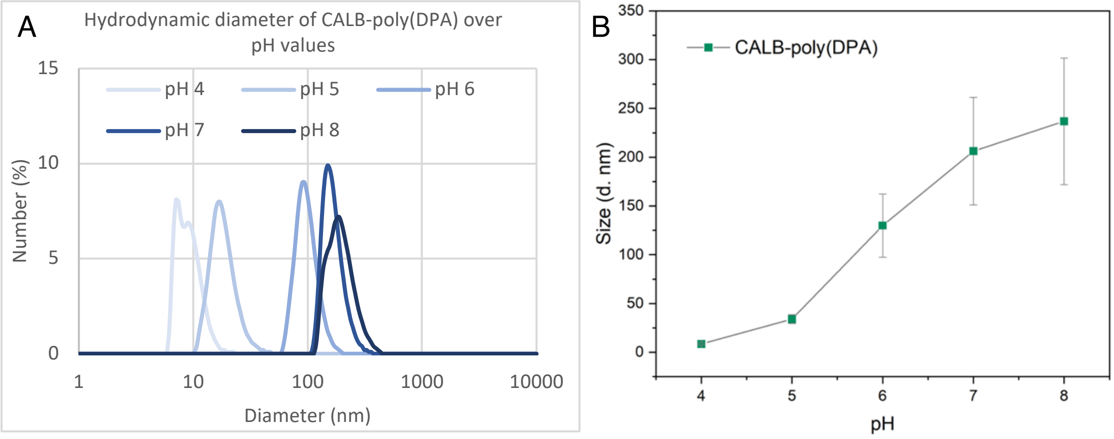


**Figure S14.** Response of CALB-poly(DPA) in 20 mM phosphate buffer on DLS. **A.** The hydrodynamic diameter distribution of CALB-poly(DPA) at pH 4, 5, 6, 7 and 8. **B.** The increase of the average hydrodynamic diameter of CALB-poly(DPA) over pH.

##### 3.7.3 Response and reversibility of the thermo-responsive TLL-poly(NIPAM)

The response and the reversibility of TLL-poly(NIPAM) was tested on UV-VIS spectrophotometer. Absorbance of TLL-poly(NIPAM) (9 µM) in 20 mM phosphate buffer solution (pH=7.4) was measured at 600 nm. TLL-poly(NIPAM) presented transmittance when was heated from 25-40 °C. The transmittance was decreased with further heating up to 42, 45 and 50 °C. The experiment was repeated successfully for multiple cycles, indicating that TLL-poly(NIPAM) maintains its responsiveness.

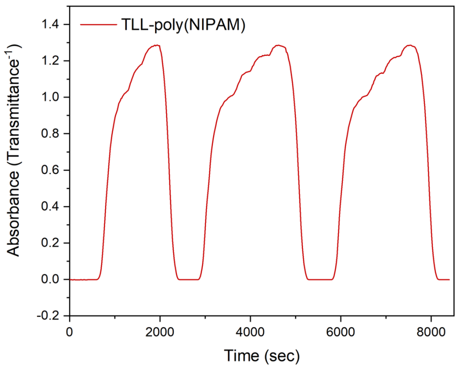


**Figure S15.** Transmittance of TLL-poly(NIPAM) in 20 mM phosphate buffer.

The responsiveness and the reversibility of TLL-poly(NIPAM) was also tested on DLS. The average hydrodynamic diameter (nm) of TLL-poly(NIPAM) was measured at 25 °C and after heating the sample at 37 °C back-and-forth five times. The average hydrodynamic diameter of TLL-poly(NIPAM) was measured from 11.76 to 15.51 nm at 25 °C and from 156.3 to 163.7 nm at 37 °C in all cases, indicating that TLL-poly(NIPAM) maintains its responsiveness.

**Table S6.** Average hydrodynamic diameter of TLL-poly(NIPAM) at 25 and 37 °C

| Cycle | Temperature (°C) | Size (d. nm) | PDI (%) |
|---|---|---|---|
| 1 | 25 | 14.14 ± 5.887 | 38.3 |
| | 37 | 163.7 ± 32.51 | 19.0 |
| 2 | 25 | 14.14 ± 8.504 | 48.8 |
| | 37 | 156.3 ± 16.07 | 10.2 |
| 3 | 25 | 15.51 ± 5.720 | 35.6 |
| | 37 | 156.3 ± 27.20 | 16.6 |
| 4 | 25 | 15.51 ± 7.431 | 44.4 |
| | 37 | 156.3 ± 23.07 | 14.5 |
| 5 | 25 | 11.76 ± 6.892 | 44.6 |
| | 37 | 163.7 ± 21.98 | 13.4 |

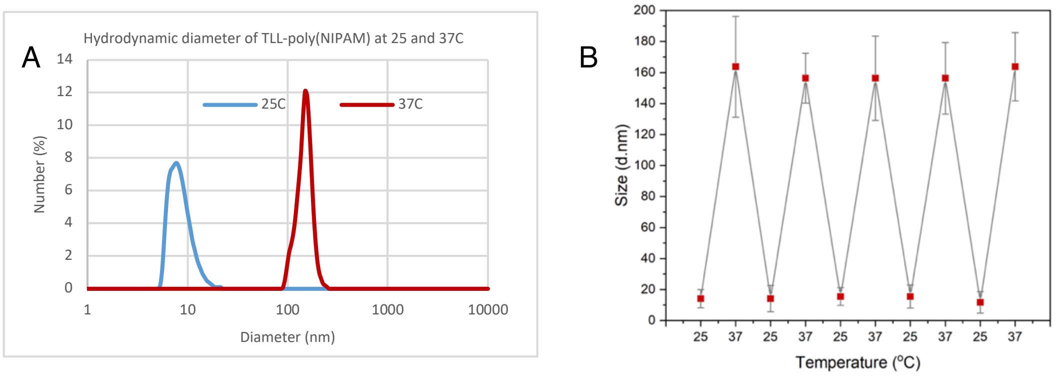


**Figure S16.** Reversibility of TLL-poly(NIPAM) in 20 mM phosphate buffer on DLS. **A.**

The hydrodynamic diameter distribution of TLL-poly(NIPAM) at 25 and 37 °C. **B**. The average hydrodynamic diameter of TLL-poly(NIPAM) was measured at 25 °C and after heating the sample at 37 °C back-and-forth five times.

The average hydrodynamic diameter (nm) of TLL-poly(NIPAM) was also measured at different temperatures (20, 25, 30, 35, 37, 40 42 and 45 °C). The average hydrodynamic diameter detected on DLS was increased after reaching the temperature of 35 °C. However, in contrary with CALB-poly(NIPAM) further increase of the temperature did not lead to higher hydrodynamic diameter values.

**Table S7.** Average hydrodynamic diameter of TLL-poly(NIPAM) heated at different temperatures

| Entry | Temperature (°C) | Size (d. Nm) | PDI (%) |
|---|---|---|---|
| 1 | 20 | 7.756 ± 1.988 | 19.4 |
| 2 | 25 | 7.406 ± 4.947 | 36.5 |
| 3 | 30 | 9.772 ± 1.355 | 12.2 |
| 4 | 35 | 216.0 ± 53.96 | 20.6 |
| 5 | 37 | 226.2 ± 53.71 | 19.7 |
| 6 | 40 | 179.5 ± 41.06 | 18.9 |
| 7 | 42 | 188.0 ± 26.57 | 12.5 |
| 8 | 45 | 179.5 ± 33.12 | 15.1 |

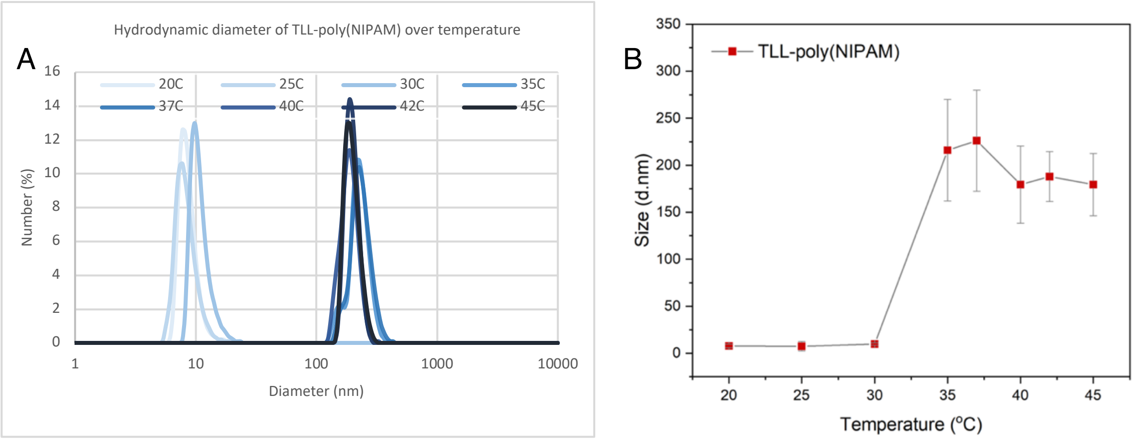


**Figure S17.** Response of TLL-poly(NIPAM) in 20 mM phosphate buffer on DLS. **A.** The hydrodynamic diameter distribution of TLL-poly(NIPAM) at 20, 25, 30, 35, 37, 40, 42 and 45 °C. **B.** The increase of the average hydrodynamic diameter of TLL-poly(NIPAM) over temperature.

The average hydrodynamic diameter (nm) of TLL-poly(NIPAM) was measured over time under constant heating at 37 °C. In contrary to CALB-poly(NIPAM) the size of TLL-poly(NIPAM) remained constant upon heating at 37 °C.

**Table S8.** Average hydrodynamic diameter of TLL-poly(NIPAM) heated at 37 °C over time

| Entry | Incubation time (min) | Size (d. nm) | PDI (%) |
|---|---|---|---|
| **1** | 5 | 158.6 ± 27.47 | 3.0 |
| **2** | 10 | 163.6 ± 35.53 | 4.7 |
| **3** | 15 | 165.3 ± 28.56 | 3.0 |
| **4** | 20 | 164.4 ± 10.18 | 0.4 |

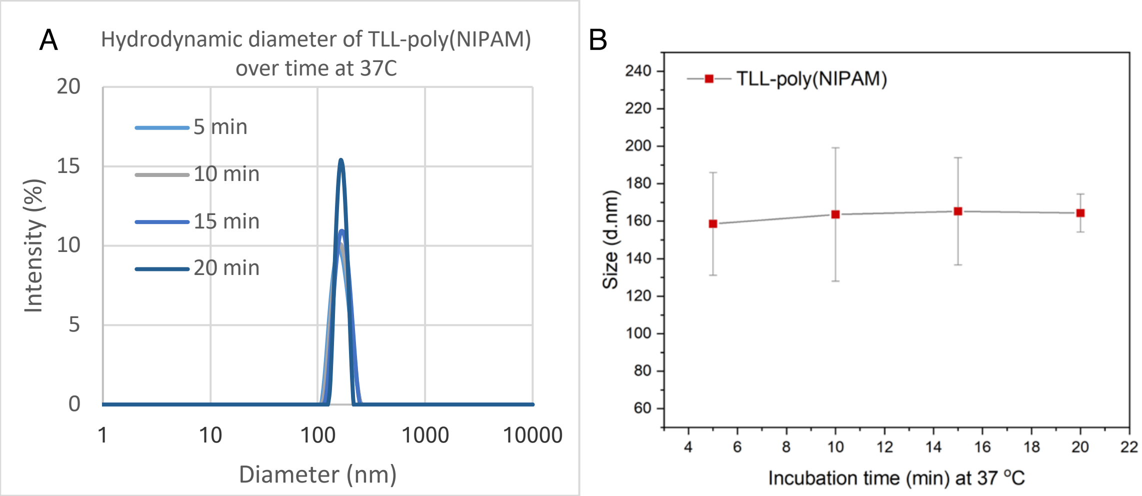


**Figure S18.** Size increase of TLL-poly(NIPAM) with constant heating at 37 °C over time measured on DLS. **A**. The hydrodynamic diameter distribution of TLL-poly(NIPAM) after 5, 10, 15 and 20 min of incubation at 37 °C. **B**. The average hydrodynamic diameter of TLL-poly(NIPAM) at each time point.

##### 3.7.4 Response of the pH-responsive TLL-poly(DPA)

The response of TLL-poly(DPA) was tested on DLS. The average hydrodynamic diameter (nm) of TLL-poly(DPA) was also measured at different pH values (4, 5, 6, 7 and 8). The average hydrodynamic diameter detected on DLS was constantly increased by pH, indicating that an assembly process is taking place at higher pH values.

**Table S9.** Average hydrodynamic diameter of TLL-poly(DPA) at different pH values

| Entry | pH value | Size (d. nm) | PDI (%) |
|---|---|---|---|
| 1 | 4 | 8.909 ± 1.300 | 14.5 |
| 2 | 5 | 68.04 ± 39.19 | 59.1 |
| 3 | 6 | 129.9 ± 31.66 | 15.5 |
| 4 | 7 | 136.1 ± 23.71 | 19.3 |
| 5 | 8 | 156.3 ± 43.27 | 21.6 |

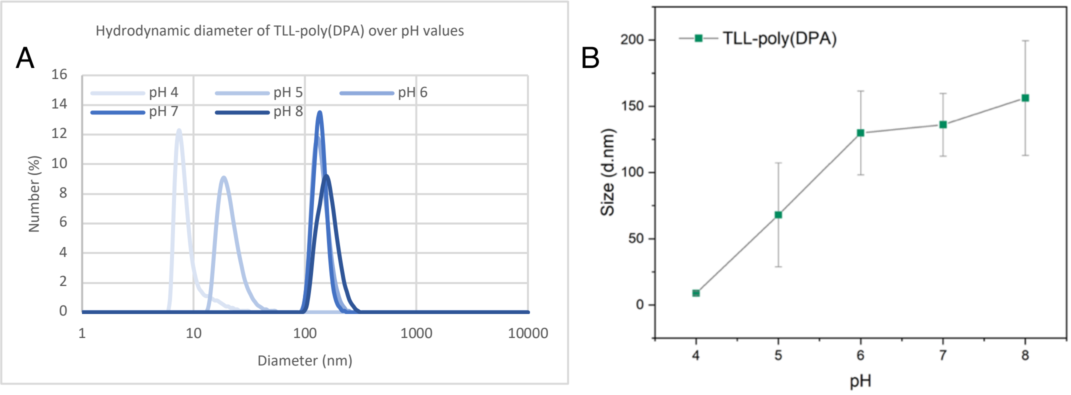


**Figure S19.** Response of TLL-poly(DPA) in 20 mM phosphate buffer on DLS. **A**. The hydrodynamic diameter distribution of TLL-poly(DPA) at pH 4, 5, 6, 7 and 8. **B**. The increase of the average hydrodynamic diameter of TLL-poly(DPA) over pH.

#### 3.8 Imaging of CALB and TLL biohybrids with total internal reflection (TIRF) microscopy

##### 3.8.1 Size distribution of CALB and TLL biohybrids

All the biohybrids were labelled with ATTO-655 NHS ester. The laser line of 640 nm was used to excite the fluorophore. Biohybrids were given 20 minutes to bind to the glass surface. Imaging was performed with an exposure time of 50 ms, 100 nm penetration depth and 300 EM gain. Each image series contained 60-120 frames of the red channel (2 mW). The image series were recorded at 4-7 fields of view for each experiment with a frame rate 2 fpm. The total integrated intensity of a vesicle labelled in its membrane (here with ATTO-655) is proportional to the square root of its diameter. The intensity readouts in combination with DLS measurements and assuming a vesicular configuration allowed us to automatically assess the size of each of the individual biohybrids and its temporal evolution in response to stimuli.

##### 3.8.1.1 Size distribution of CALB coated poly(styrene)

The degree of labelling was measured on Nanodrop and found to be 0.18 +/- 0.03 %. The number of particles detected in 6 fields of view was 300. Due to the spherical structure of CALB coated poly(styrene) particles, the diameter is analogous to the square root of the intensity,[3] allowing us to convert the intensity distribution to a size distribution using the mean diameter value found in DLS measurements. The histograms are fitted with the LogNormal fit. The mu values and the sigma of the LogNormal fit are displayed.

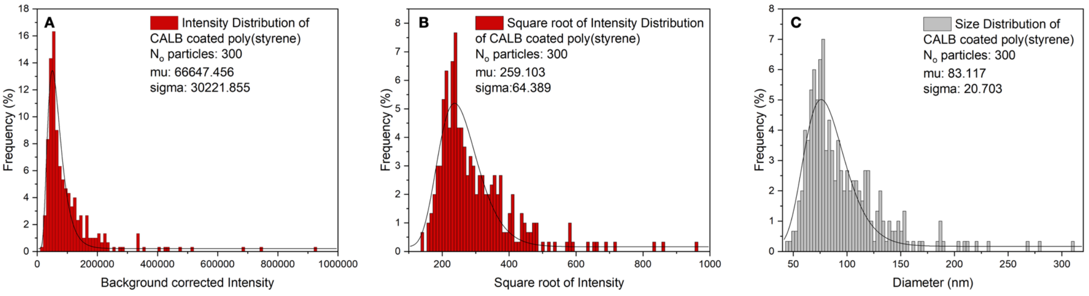


**Figure S20. A.** Distribution of background corrected intensities of CALB coated poly(styrene). **B**. Distribution of the square root of intensities of CALB coated poly(styrene). **C**. Distribution of diameter values of CALB coated poly(styrene).

##### 3.8.1.2 Size distribution of the temperature responsive CALB-poly(NIPAM)

The degree of labelling was measured on Nanodrop and found to be 0.32 +/- 0.13%. The number of particles detected in 5 fields of view was 473 imaging at 25 °C. The number of particles detected in 6 fields of view was 1117 imaging at 37 °C. CALB-poly(NIPAM) forms spherical assemblies at 37 °C, whereas exists at the monomeric form at 25 °C. Therefore, only the intensity distribution at 37 °C was converted to size distribution. The histograms are fitted with the LogNormal fit. The mu values and the sigma of the LogNormal fit are displayed.

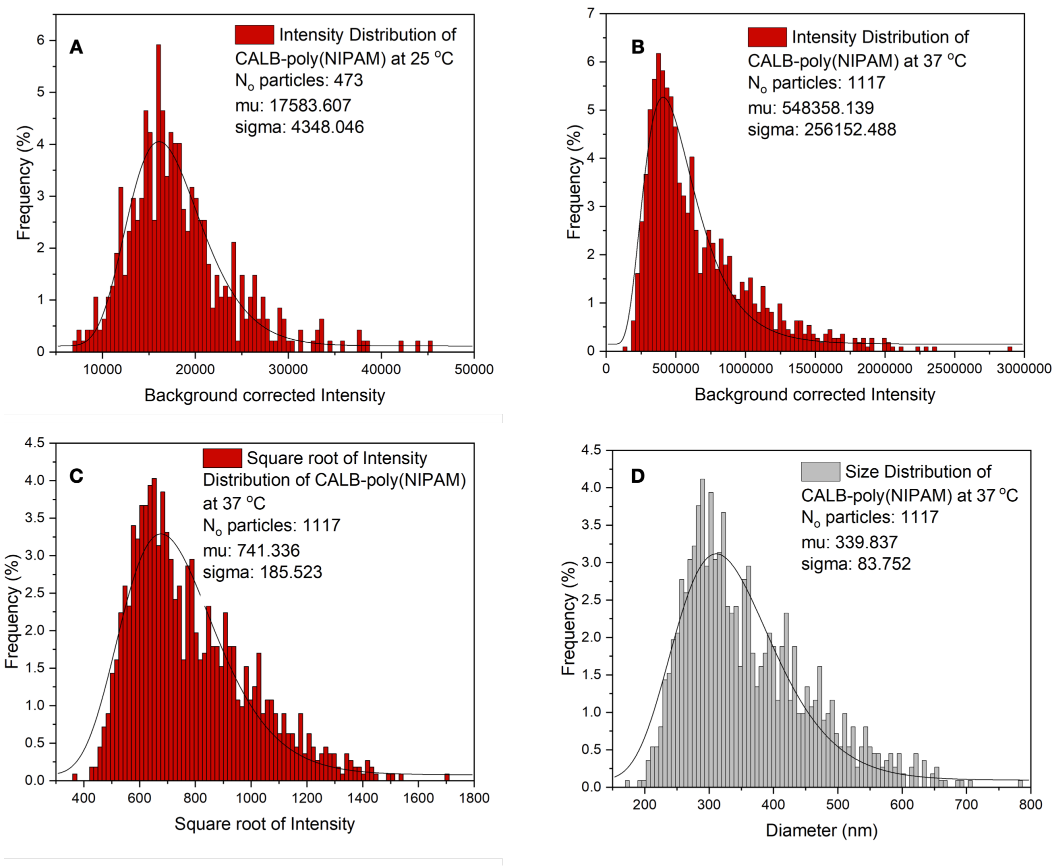


**Figure S21. A.** Distribution of background corrected intensities of CALB-poly(NIPAM) at 25 °C. **B**. Distribution of background corrected intensities of CALB-poly(NIPAM) at 37 °C. **C**. Distribution of the square root of intensities of CALB-poly(NIPAM) at 37 °C. **D**. Distribution of diameter values of CALB-poly(NIPAM) at 37 °C.

##### 3.8.1.3 Size distribution of the pH responsive CALB-poly(DPA)

The degree of labelling was measured on Nanodrop and found to be 0.26 +/- 0.06 %. The number of particles detected in 4 fields of view was 341 imaging at pH 5.0. The number of particles detected in 4 fields of view was 2474 imaging at pH 8.2. At these pH values CALB-poly(DPA) forms spherical assemblies of different sizes. The histograms are fitted with the LogNormal fit. The mu values and the sigma of the LogNormal fit are displayed.

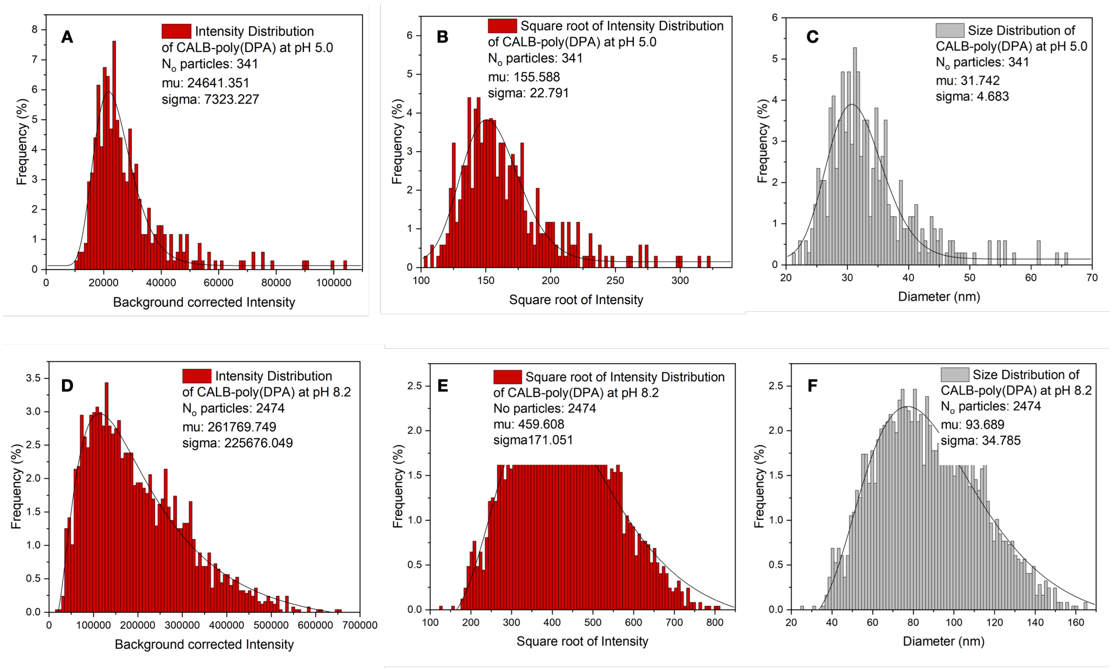


**Figure S22. A.** Distribution of background corrected Intensities of CALB-poly(DPA) at pH 5.0. **B.** Distribution of the square root of Intensities of CALB-poly(DPA) at pH 5.0. **C**. Distribution of diameter values of CALB-poly(DPA) at pH 5.0. **D.** Distribution of background corrected Intensities of CALB-poly(DPA) at pH 8.2. **E**. Distribution of the square root of Intensities of CALB-poly(DPA) at pH 8.2. **F**. Distribution of diameter values of CALB-poly(DPA) at pH 8.2.

##### 3.8.1.4 Size distribution of TLL-poly(styrene)

The degree of labelling was measured on Nanodrop and found to be 0.10 +/- 0.02%. The number of particles detected in 6 fields of view was 234. TLL-poly(styrene) forms spherical structures, therefore, the intensity distribution was converted to size distribution. The histograms are fitted with the LogNormal fit. The mu values and the sigma of the LogNormal fit are displayed.

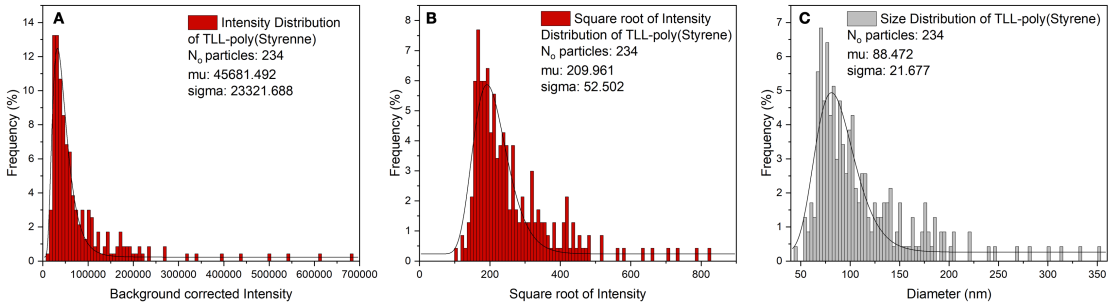


**Figure S23. A.** Distribution of background corrected Intensities of TLL-poly(styrene).

**B**. Distribution of the square root of Intensities of TLL-poly(styrene). **C**. Distribution of diameter values of TLL- poly(styrene).

##### 3.8.1.5 Size distribution of the temperature responsive TLL-poly(NIPAM)

The degree of labelling was measured on Nanodrop and found to be 0.16 +/- 0.06%. The number of particles detected in 6 fields of view was 316 imaging at 25 °C. The number of particles detected in 6 fields of view was 1065 imaging at 37 °C. TLL-poly(NIPAM) forms spherical assemblies at 37 °C, whereas exists at the monomeric form at 25 °C. Therefore, only the intensity distribution at 37 °C was converted to size distribution. The histograms are fitted with the LogNormal fit. The mu values and the sigma of the LogNormal fit are displayed.

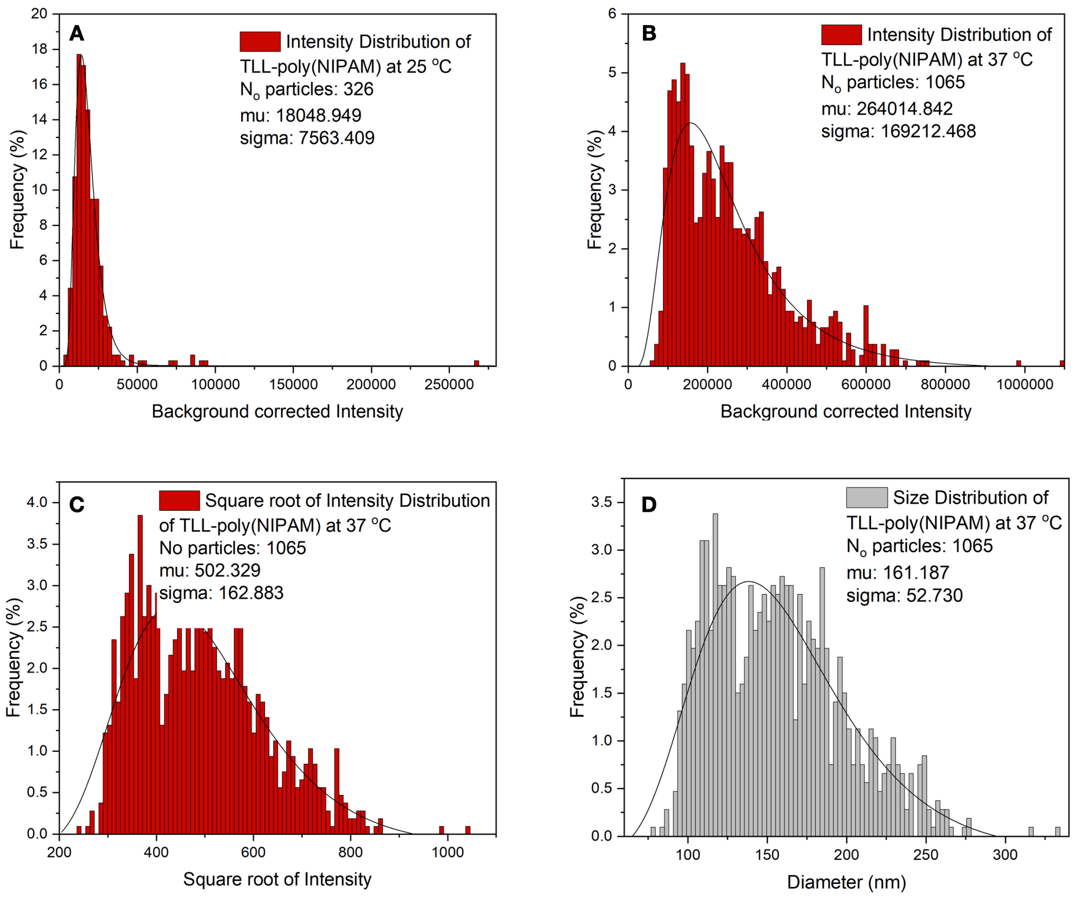


**Figure S24. A.** Distribution of background corrected intensities of TLL-poly(NIPAM) at 25 °C. **B**. Distribution of background corrected intensities of TLL-poly(NIPAM) at 37 °C. **C**. Distribution of the square root of intensities of TLL-poly(NIPAM) at 37 °C. **D**. Distribution of diameter values of TLL-poly(NIPAM) at 37 °C.

##### 3.8.1.6 Size distribution of the pH responsive TLL-poly(DPA)

The degree of labelling was measured on Nanodrop and found to be 0.20 +/- 0.17%.

The number of particles detected in 7 fields of view was 506 imaging at pH 5.0. The number of particles detected in 6 fields of view was 1203 imaging at pH 8.2. At these pH values TLL-poly(DPA) forms spherical assemblies of different sizes. The histograms are fitted with the LogNormal fit. The mu values and the sigma of the LogNormal fit are displayed.

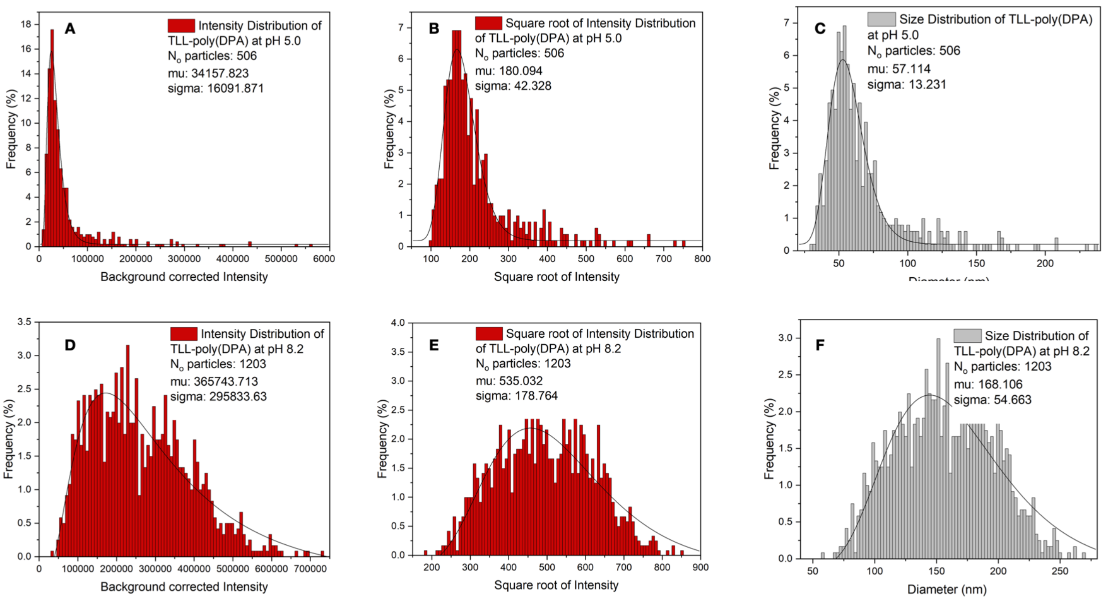


**Figure S25. A.** Distribution of background corrected intensities of TLL-poly(DPA) at pH 5.0. **B**. Distribution of the square root of intensities of TLL-poly(DPA) at pH 5.0. **C.** Distribution of diameter values ofTLL-poly(DPA) at pH 5.0. **D**. Distribution of background corrected intensities of TLL-poly(DPA) at pH 8.2. **E**. Distribution of the square root of intensities of TLL-poly(DPA) at pH 8.2. **F.** Distribution of diameter values of TLL-poly(DPA) at pH 8.2.

##### 3.8.2 Single enzyme calibration curve – Calculation of N$_o$ enzymes in biohybrids

Native TLL labelled with ATTO-655 NHS ester was imaged using the laser line of 640 nm to excite the fluorophore (2 mW). Imaging was performed with an exposure time of 50, 100, 300, 400 and 500 nm, 100 nm penetration depth and 300 EM gain. The formula for linear regression was ***f(x)=237.54x + 4663.70***.

| Exposure time (ms) | Mean intensity | S.D. |
|---|---|---|
| 50 | 10828.511 | 1978.491 |
| 100 | 29173.501 | 4825.397 |
| 300 | 80185.236 | 14567.402 |
| 400 | 113843.450 | 23782.563 |
| 500 | 109967.791 | 21267.652 |

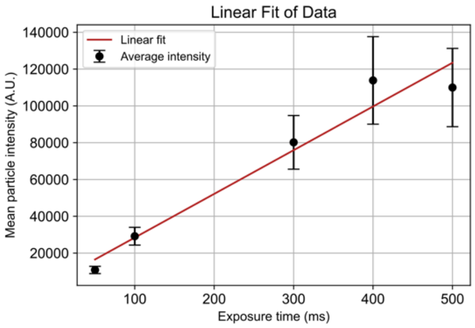


**Figure S26.** Mean intensity values of the intensity distribution of native TLL imaged with 5 different exposure times and the linear fit of the data based on three technical replicates where error is calculated as the standard deviation of the mean of a normal fit of the intensity distribution.

Quantitative image analysis allowed quantification of the total number of enzymes in each type of biohybrid under varying temperature and pH conditions. Our calculations were based on our previous methodologies[3,4] and considering the degree of labelling (DOL) using the equation (i) we found that for 50 ms exposure time at 25 $^{o}$C CALB- and TLL-poly(NIPAM) exist at the monomeric form (similar intensity with single enzymes), whereas at 37 $^{o}$C these biohybrids form assemblies consisted of ~11k enzymes. CALB-poly(DPA) forms assemblies with varying sizes depending on the pH with the number of enzymes to vary from 559 to almost 6k from pH 5 to pH 8 respectively. Similarly, for TLL-poly(DPA) at pH 5 the biohybrid forms small assemblies of 1K enzymes and higher order assemblies of 11k enzymes at pH 8.

$$Number\ of\ enzymes = \frac{Mean\ intensity\ of\ the\ biohybrid}{Mean\ intensity\ of\ the\ single\ enzyme}\ x\ \frac{100}{DOL}\ (i)$$

**Table S10.** Number of enzymes in each biohybrid calculations

| Biohybrid | Enzyme concentration (µM) | Dye concentration (µM) | Degree of labelling (%) | DLS (d.nm) | TIRF (d.nm) | Median Intensity (mu) | Number of enzymes |
|---|---|---|---|---|---|---|---|
| CALB coated poly(styrene) | 45 | 0,083 +/- 0,015 | 0.18 +/- 0.03 | 83.55 | 83.117 | 66647.456 | 2145.657+/- 362.616 |
| CALB-poly(NIPAM) at 25 $^{o}$C | 20 | 0,063 +/- 0,025 | 0.32 +/- 0.13 | 11.54 | - | 17583.607 | Mn |
| CALB-poly(NIPAM) at 37 $^{o}$C | 20 | 0,063 +/- 0,025 | 0.32 +/-0.13 | 345.3 | 339.837 | 548358.139 | 11029.844 +/- 4870.413 |
| CALB-poly(DPA) at pH 5.0 | 10 | 0,026 +/- 0,006 | 0.26 +/- 0.06 | 31.69 | 31.742 | 24641.351 | 559.053 +/- 132.504 |
| CALB-poly(DPA) at pH 8.2 | 10 | 0,026 +/- 0,006 | 0.26 +/- 0.06 | 87.71 | 93.689 | 261769.749 | 5940.486 +/- 1407.989 |

| | | | | | | | |
|---|---|---|---|---|---|---|---|
| TLL-poly(styrene) | 30 | 0,026 +/- 0,006 | 0.1 +/- 0.02 | 89.27 | 88.472 | 45681.492 | 2601.306 +/- 530.795 |
| TLL-poly(NIPAM) at 25 $^{o}$C | 10 | 0,016 +/- 0,006 | 0.16 +/- 0.06 | 18.26 | - | 18048.949 | Mn |
| TLL-poly(NIPAM) at 37 $^{o}$C | 10 | 0,016 +/- 0,006 | 0.16 +/- 0.06 | 163.0 | 161.187 | 264014.842 | 10139.864 +/- 4081.886 |
| TLL-poly(DPA) at pH 5.0 | 10 | 0,02 +/- 0,007 | 0.2 +/- 0.07 | 56.83 | 57.114 | 34157.823 | 1034.588 +/- 385.152 |
| TLL-poly(DPA) at pH 8.2 | 10 | 0,02 +/- 0,007 | 0.2 +/- 0.07 | 142.4 | 168.106 | 365743.713 | 11074.959 +/- 4122.937 |

##### 3.8.3 Response of CALB and TLL biohybrids at single particle level

###### 3.8.3.1 Intensity increase of the temperature responsive CALB-poly(NIPAM) upon heating at 37 $^{o}$C for 1 hour.

CALB-poly(NIPAM) labelled with ATTO-655 NHS ester was incubated at 37 $^{o}$C for 30 min to self-assemble. Then, it was imaged while heated 37 $^{o}$C for 1 hour using the laser line of 640 nm, an exposure time of 50 ms, 100 nm penetration depth and 300 EM gain. Images were recorded upon heating with a frame rate of 2 fpm.

**Table S11.** Background corrected mean intensities over time under constant heating at 37 $^{o}$C

| Entry | Time (min) | Mean Intensity values | S. D. |
|---|---|---|---|
| **1** | 0 | 743960.605 | 415778.949 |
| **2** | 5 | 866549.249 | 455796.536 |
| **3** | 10 | 1006123.540 | 541698.747 |
| **4** | 15 | 1108576.182 | 601681.269 |
| **5** | 20 | 1264763.032 | 688406.092 |
| **6** | 25 | 1302407.071 | 781104.903 |
| **7** | 30 | 1385089.511 | 820940.557 |
| **8** | 35 | 1484133.665 | 874513.930 |
| **9** | 40 | 1593239.397 | 902959.837 |
| **10** | 45 | 1610256.553 | 948952.697 |
| **11** | 50 | 1654756.218 | 993247.226 |
| **12** | 55 | 1653293.127 | 974130.7467 |
| **13** | 60 | 1704468.851 | 1028895.946 |

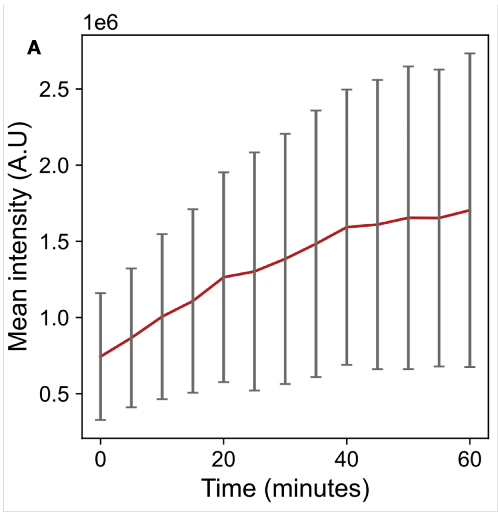


**Figure S27.** **A**. Mean background corrected intensities with the standard deviation of the mean value of every 5 min (10 frames) of a series of images recorded upon heating CALB-poly(NIPAM) at 37 °C for 1 hour. Error bars correspond to the standard deviation of the mean intensity values for all the CALB-poly(NIPAM) particles every 5 min (10 frames).

##### 3.8.3.2 Disassembling of the temperature responsive CALB- and TLL-poly(NIPAM) from 37 to 25 °C on TIRF

CALB- and TLL-poly(NIPAM) labelled with ATTO-655 NHS ester was incubated at 37 °C for 30 min to self-assemble. Then, it was imaged at 37 °C using the laser line of 640 nm, an exposure time of 50 ms, 100 nm penetration depth and 300 EM gain. The number of particles detected in 6 fields of view was 283 and 2455 and at 37 °C for CALB- and TLL-poly(NIPAM) respectively. The sample was left for 2 hours to be cooled down to 25 °C and imaged again at the same fields of view. The number of particles detected in 6 fields of view was 595 and 426 at 25 °C for CALB- and TLL-poly(NIPAM) respectively. The histograms are fitted with the LogNormal fit. The mean values and the sigma of the LogNormal fit are displayed.

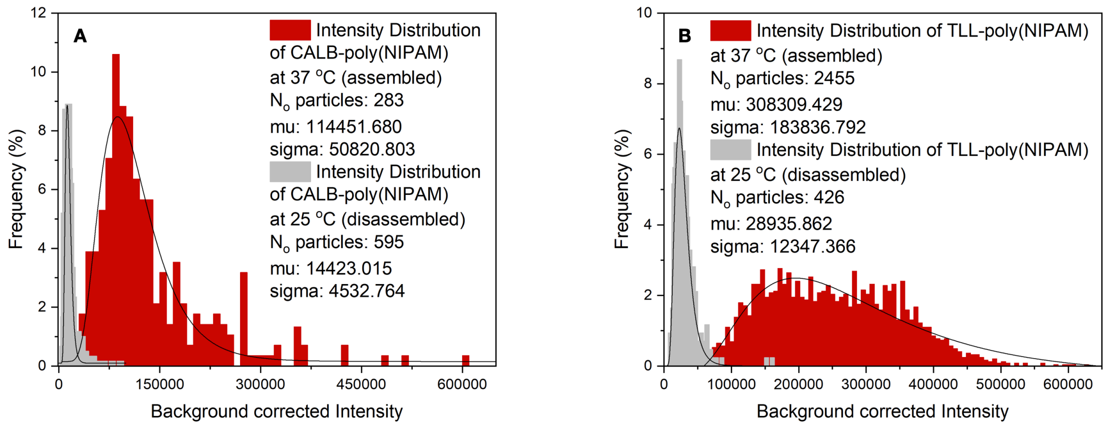


**Figure S28.** **A**. Distributions of background corrected intensities of CALB-poly(NIPAM) at 37 °C and after the sample was cooled down to 25 °C. **B**. Distributions of background corrected intensities of TLL-poly(NIPAM) at 37 °C and after the sample was cooled down to 25 °C.

#### 3.9 Catalytic activity of CALB and TLL biohybrids (bulk measurements)

##### 3.9.1 Catalytic activity of native CALB and CALB biohybrids over CFDA at 37 °C pH 7.4

1 mg of 5-(6)-carboxyfluorescein diacetate (CFDA) was dissolved in 250 µL DMSO to form a stock solution which was fractionated and stored at -20 °C. 15 µL of a 0.31 mM CALB solution were diluted with 50 mM phosphate buffer pH 7.4 (2 mL) to form a 2.33 µM solution. The reaction was initiated by the addition of the 25 µL of the CFDA solution to the CALB solution and the esterase-like activity of CALB was monitored by UV at 453 nm at 37 °C. The ability of CALB-biohybrids to hydrolyse CFDA was also tested following the same protocol. A blank experiment was performed using the same protocol in the absence of the enzyme. All reactions were performed in triplicates.

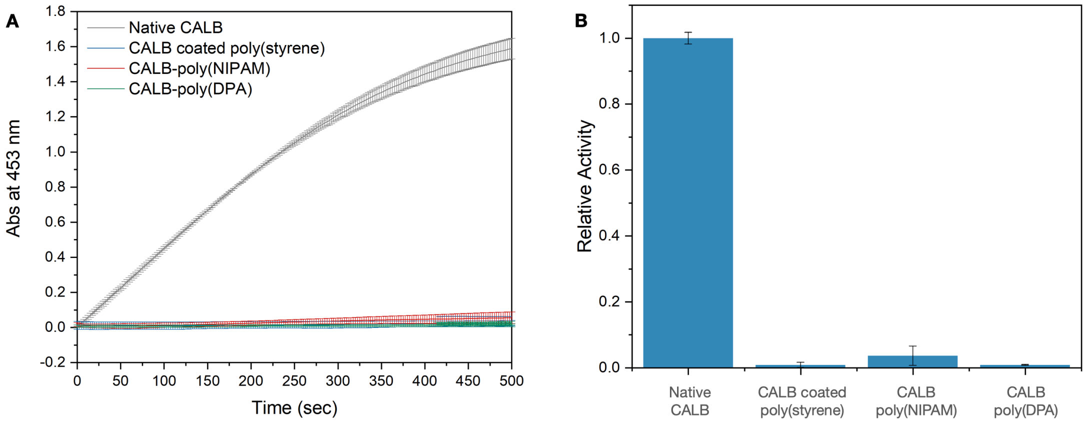


**Figure S29.** **A**. Esterase-like activity of native CALB and CALB biohybrids at 37 °C on CFDA substrate. **B.** Relative activity of CALB biohybrids to native CALB at 37 °C.

##### 3.9.2 Activity of native CALB and the temperature responsive CALB-poly(NIPAM) at a series of temperatures

The esterase-like activity of the native CALB and thermo-responsive CALB-poly(NIPAM) was tested at 20, 25 and 37 °C following the same protocol described above. For the temperature responsive CALB-poly(NIPAM) the sample was diluted in the 50 mM phosphate buffer pH 8.2 (2 mL), incubated at each temperature for 10 min to self-assemble, followed by the addition of the CFDA solution. All reactions were performed in triplicates.

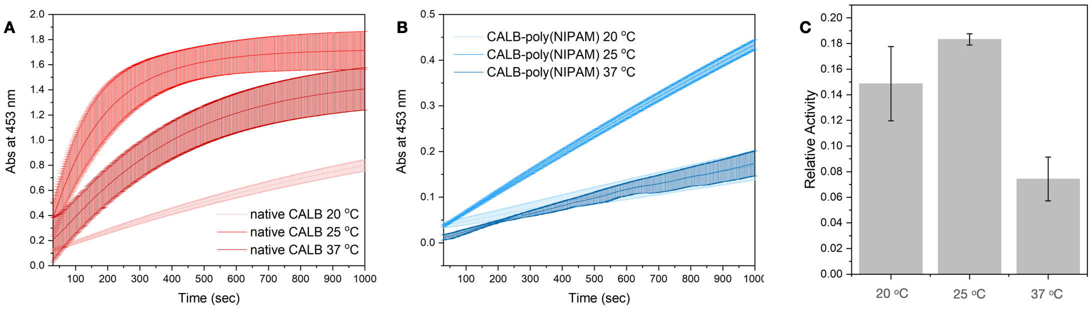


**Figure S30. A**. Esterase-like activity of native CALB at 20, 25 and 37 °C on CFDA substrate. **B**. Esterase-like activity of CALB-poly(NIPAM) at 20, 25 and 37 °C on CFDA substrate. **C.** Relative activity of CALB-poly(NIPAM) to native CALB at each temperature.

##### 3.9.3 Catalytic activity of native TLL, TLL-poly(styrene) and TLL-poly(DPA) over CFDA at 37.5 °C pH 8.2

1 mg of 5-(6)-carboxyfluorescein diacetate (CFDA) was dissolved in 250 µL DMSO to form a stock solution which was fractionated and stored at -20 °C. 9 µL of a 0.52 mM TLL solution were diluted with 50 mM phosphate buffer pH 8.2 (2 mL) to form a 2.34 µM solution. The reaction was initiated by the addition of the 25 µL of the CFDA solution to the TLL solution and the esterase-like activity of TLL was monitored by UV at 453 nm at 37.5 °C. The ability of TLL-Br and TLL biohybrids to hydrolyze CFDA was also tested following the same protocol. A blank experiment was performed using the same protocol in the absence of the enzyme. All reactions were performed in triplicates.

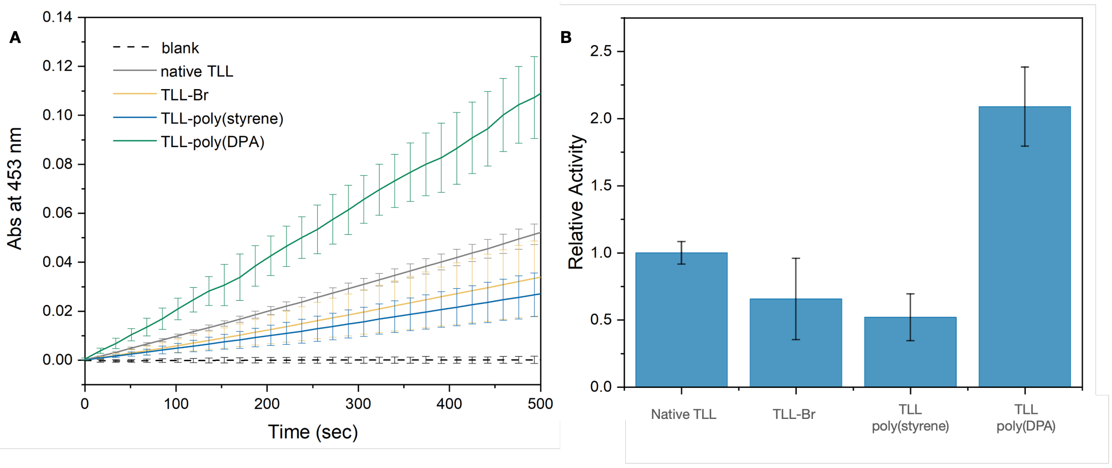


**Figure S31.** **A**. Esterase-like activity of native TLL, TLL-Br and TLL biohybrids at 37.5 °C on CFDA substrate. **B.** Relative activity of TLL biohybrids to native TLL at 37.5 °C.

##### 3.9.4 Catalytic activity of native TLL and the temperature responsive TLL-poly(NIPAM) at a series of temperatures

The esterase-like activity of the native TLL and thermo-responsive TLL-poly(NIPAM) was tested at 25, 37.5, 40, 42 and 45 °C following the same protocol described above. For the temperature responsive TLL-poly(NIPAM) the sample was diluted in the 50 mM phosphate buffer pH 8.2 (2 mL), incubated at each temperature for 10 min to self-assemble, followed by the addition of the CFDA solution. All reactions were performed in triplicates. At all the tested temperatures TLL-poly(NIPAM) presented higher activity compared to the native TLL. The higher activation of native TLL compared to TLL-poly(NIPAM) was observed at 37.5 and 40 °C,

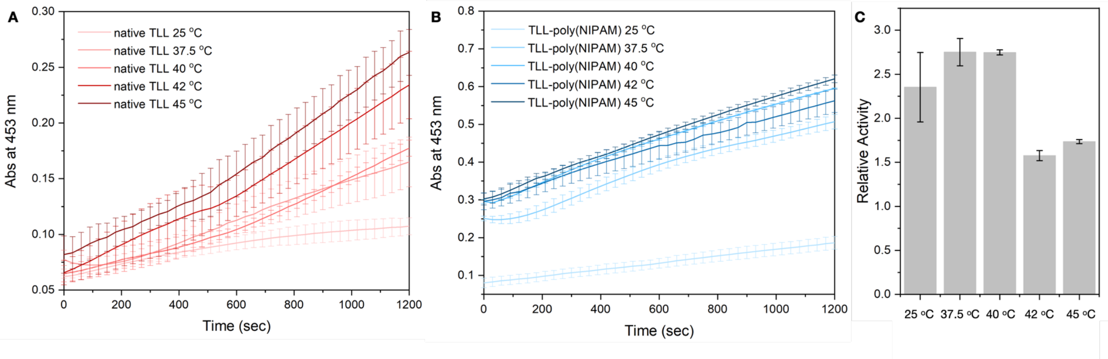


**Figure S32.** **A**. Esterase-like activity of native TLL at 25, 37.5, 40, 42 and 45 °C on CFDA substrate. **B**. Esterase-like activity of TLL-poly(NIPAM) at 25, 37.5, 40, 42 and 45 °C on CFDA substrate. **C.** Relative activity of TLL-poly(NIPAM) to native TLL at each temperature.

#### 3.10 Catalytic activity of CALB and TLL biohybrids at single particle level

The catalytic activity of CALB and TLL biohybrids was also studied at single particle level using total internal reflection (TIRF) microscopy.

All the biohybrids were labelled with ATTO-655 NHS ester and could be detected in the red channel. CFDA was used as the substrate. The hydrolysis product ($CF^-$) is a fluorescent compound that has an emission peak at 517 nm. Therefore, when the hydrolysis products were attached on the surface of the biohybrids we could detect signal in the blue channel. The laser lines of 640 nm and 488 were used to excite the fluorophore and the hydrolysis product respectively. Biohybrids in 20 mM phosphate buffer pH 8.2 (10 nM in the final solution) were given 20 minutes to bind to the glass surface, followed by the addition of CFDA solution (30 nM in the final solution). Then, Imaging was performed with an exposure time of 50 ms, 100 nm penetration depth and 300 EM gain. Each image series contained 60 frames of the red channel (2 mW) and 60 frames of the blue channel (1.3 mW). The image series were recorded at 5-6 fields of view for each experiment with a frame rate 2 fpm. Image series under the same experimental condition were recorded in the absence of CFDA as a control experiment.

##### 3.10.1 Catalytic activity of CALB biohybrid at singe particle level

**Table S11** Background corrected mean intensity in the hydrolysis product channel for the CALB biohybrids

| Biohybrid | Background corrected intensity in the presence of CFDA | Background corrected intensity in the absence of CFDA (control) | *ΔI* |
|---|---|---|---|
| CALB coated poly(styrene | 3776.979 | 1271.629 | 2505.35 |
| CALB-poly(NIPAM) at 25 °C | 358.659 | 337.329 | 21.33 |
| CALB-poly(NIPAM) at 37 °C | 37110.275 | 8990.308 | 28119.967 |
| CALB-poly(DPA) | 339.579 | 314.801 | 24.778 |

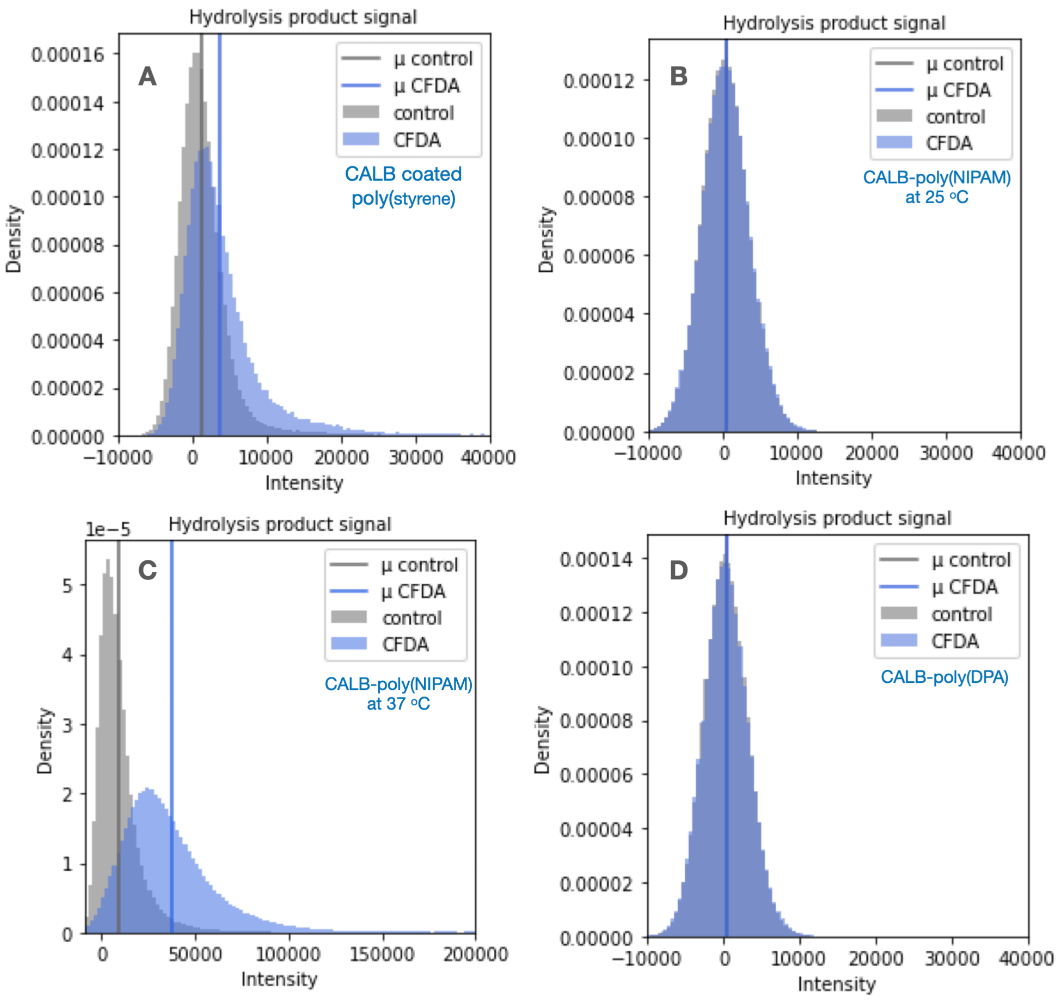


**Figure S33.** **A**. Distribution of background corrected intensities in the blue channel of the CALB coated poly(styrene) in the absence (control) and in the presence of CFDA. **B**. Distribution of background corrected intensities in the blue channel of the CALB-poly(NIPAM) at 25 °C in the absence (control) and in the presence of CFDA. **C.** Distribution of background corrected intensities in the blue channel of the CALB-poly(NIPAM) at 37 °C in the absence (control) and in the presence of CFDA. **D**. Distribution of background corrected intensities in the blue channel of the CALB-poly(DPA) in the absence (control) and in the presence of CFDA.

##### 3.10.2 Catalytic activity of TLL biohybrid at singe particle level

**Table S12** Background corrected mean intensity in the hydrolysis product channel for the TLL biohybrids

| Biohybrid | Background corrected intensity in the presence of CFDA | Background corrected intensity in the absence of CFDA (control) | *ΔI* |
|---|---|---|---|
| TLL- poly(styrene) | 1471.472 | 1390.697 | 80.778 |
| TLL-poly(NIPAM) at 25 °C | 794.247 | 790.119 | 4.128 |
| TLL-poly(NIPAM) at 37 °C | 2882.694 | 1434.073 | 1448.621 |
| TLL-poly(DPA) | 1471.472 | 1192.246 | 279.226 |

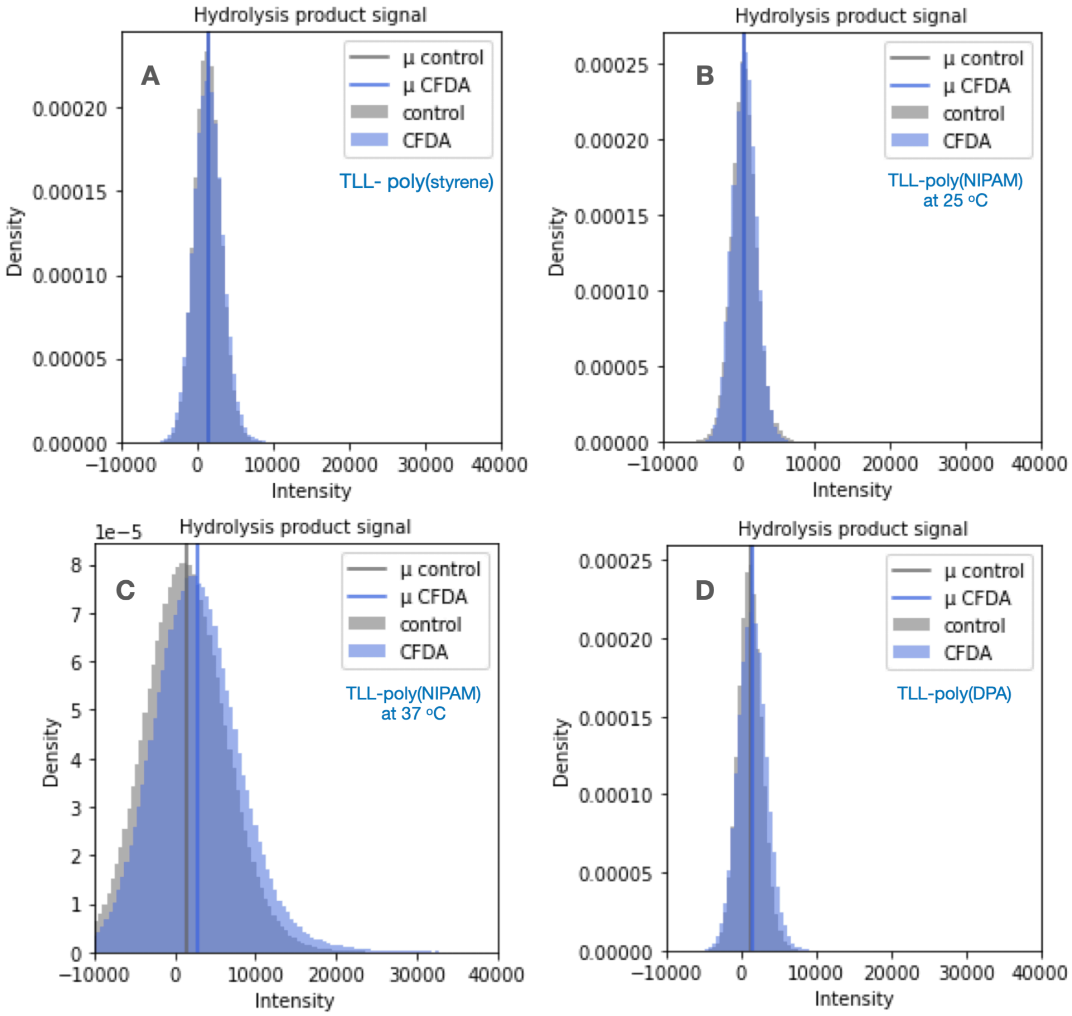


**Figure S34.** **A**. Distribution of background corrected intensities in the blue channel of the TLL-poly(styrene) in the absence (control) and in the presence of CFDA. **B**. Distribution of background corrected intensities in the blue channel of the TLL-poly(NIPAM) at 25 °C in the absence (control) and in the presence of CFDA. **C.** Distribution of background corrected intensities in the blue channel of the TLL-poly(NIPAM) at 37 °C in the absence (control) and in the presence of CFDA. **D**. Distribution of background corrected intensities in the blue channel of the TLL-poly(DPA) in the absence (control) and in the presence of CFDA.

#### 3.11 Number of turnovers per enzyme in CALB-poly(NIPAM) at 37 °C

##### 3.11.1 Hydrolysis product calibration curve

Surface preparation:

Glass slides 76 x 26 nm were dried with a nitrogen flow and activated using a plasma cleaner for 1 minute at a pressure between 300 mtorr to 400 mtorr. Immediately after activation the slides were attached to the glass slides in order to make flow cells. Each of the six champers was passivated with 80 µL of a PLL-g-PEG (1 mg/mL in HEPES buffer pH 5.5) and PLL-g-PEG-biotin (1 mg/mL in HEPES buffer pH 5.5) mixture in the ratio 100:1. The surface was incubated for 30 min. Excess of PLL-g-PEG mixture was removed by flushing five times with 50 µL HEPES buffer pH 5.5. 80 µL of the neutravidin solution (1 mg/mL in HEPES buffer pH 5.5) was added in each well. The surface was incubated for 15 min. Each well was washed with 20 mM PB buffer pH 8.2 at least 5 times before the sample addition.

Biotinylated fluorescein (hydrolysis product) was tethered in the neutravidin surface and was imaged using the laser line of 488 nm to excite the hydrolysis product (1.3 mW). Imaging was performed with an exposure time of 50, 200, 400 and 500 nm, 100 nm penetration depth and 300 EM gain. The formula for linear regression was ***f(x)=156.80x + 12670.83***.

| Exposure time (ms) | Mean intensity | S.D. |
|---|---|---|
| 50 | 20268.089 | 4451.698 |
| 200 | 43968.904 | 9834.757 |
| 400 | 76670.918 | 13624.317 |
| 500 | 90097.275 | 16916.345 |

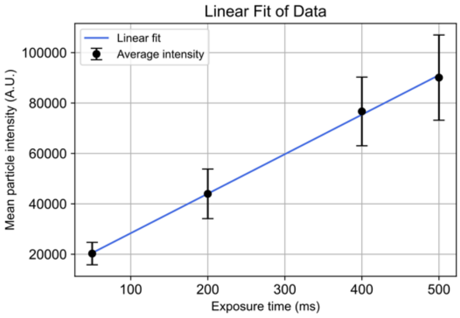


**Figure S35.** Calibration curve of the turnovers per a single lipase imaged with 4 different exposure times and the linear fit of the data, based on three technical replicates where error is calculated as the standard deviation of the mean of a normal fit of the intensity distribution.

Since CALB-poly(NIPAM) is a polydisperse sample, spherical structures of different size have a different catalytic activity outcome. According to our calibration curve, for 50 ms exposure time the hydrolysis product has a mean intensity f(50)=20510.83. Based on this we calculated the number of turnovers per biohybrid size and per enzyme in each biohybrid size for the CALB-poly(NIPAM) at 37 °C. Biohybrids with larger sizes (higher number of enzymes) have higher catalytic outcome as expected. However, in a single enzyme level, biohybrids of smaller sizes (lower number of enzymes) are composed of enzymes with higher catalytic outcome.

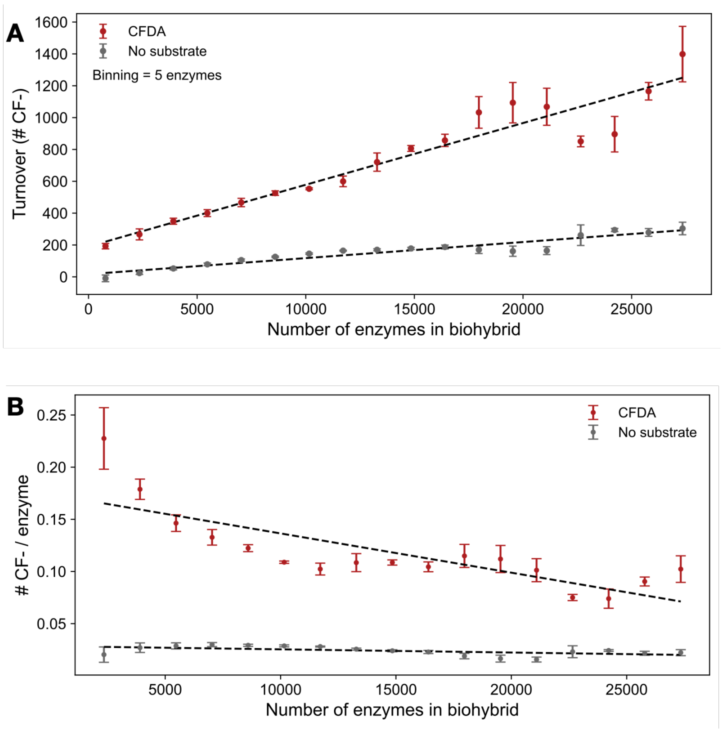


**Figure S36. A.** Number of turnovers per CALB-poly(NIPAM) size at 37 °C. estimated by the turnovers per a single lipase calculated by the calibration curve presented in figure S36 and the number of enzymes in CALB-poly(NIPAM) size at 37 °C, based on six technical replicates where the error shows the standard deviation between replicates. **B**. Number of turnovers per enzyme in each CALB-poly(NIPAM) size at 37 °C based on six technical replicates where the error shows the standard deviation between replicates.

#### 3.12 Stability of CALB and TLL biohybrids

##### 3.12.1 Stability of native CALB and CALB biohybrids after 1 year of storage at 4 °C.

The activity of native CALB and CALB biohybrids was measured at 25 °C immediately and 1 year after preparation and stored at 4 °C. The fresh and the 1-year-old samples presented similar ability to hydrolyze CFDA indicating the high stability of the CALB biohybrids. All the CALB biohybrids presented higher stability compared to the native CALB. Native CALB lost almost 40% of the catalytic activity after 1 year of storage. CALB coated poly(styrene), CALB-poly(NIPAM) and CALB-poly(DPA) lost almost 10, 5 and 20% of their catalytic activity after 1 year of storage respectively.

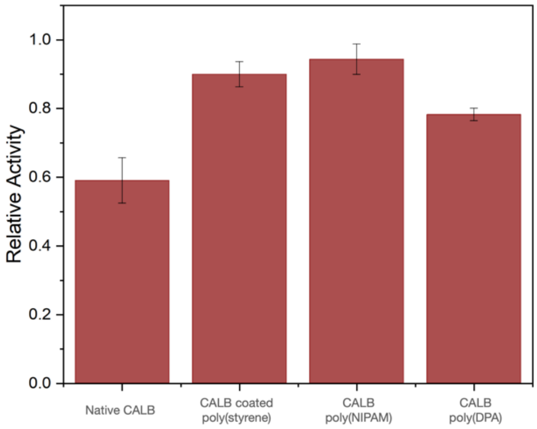


| Biohybrid | Relative activity | S.D. |
|---|---|---|
| Native CALB | 0.59018 | 0.06594 |
| CALB coated poly(styrene) | 0.89979 | 0.03665 |
| CALB-poly(NIPAM) | 0.94354 | 0.04417 |
| CALB-poly(DPA) | 0.78236 | 0.01816 |

**Figure S37.** Relative activity of the 1-year-old sample to the fresh sample for the native CALB and each biohybrid.

##### 3.12.2 Stability of native TLL and TLL biohybrids after 1 year of storage at 4 °C.

The activity at 37.5 °C of native TLL and TLL biohybrids was also measured 2 and 12 months after preparation and storage at 4 °C using an Id-3 spectraMax plate reader. A clear standard 96-well plates were used to measure absorbance at 453 nm. All reactions were performed in triplicates. The 2- and 12-month-old TLL-poly(styrene) presented similar ability to hydrolyse CFDA, whereas TLL-poly(NIPAM) and TLL-poly(DPA) presented similar stability to native TLL retaining part of their catalytic activity.

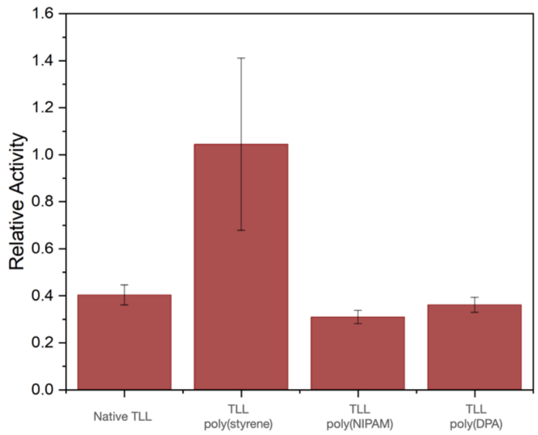


| Biohybrid | Relative activity | S.D. |
|---|---|---|
| Native TLL | 0.40217 | 0.04246 |
| TLL-poly(styrene) | 1.04406 | 0.3666 |
| TLL-poly(NIPAM) | 0.30812 | 0.02795 |
| TLL-poly(DPA) | 0.36002 | 0.03203 |

**Figure S38.** Relative activity of the 12-month-old sample to the 2-month-old sample for the native TLL and each biohybrid.

#### 3.13 CALB batch variation

The CALB batch variation was studied to detect the consistency between different production batches. The catalytic activity of three CALB batches and the CALB-poly(NIPAM) produced from these batches was tested over CFDA substate at 20 and 37 °C, following the experimental procedure described in paragraph 3.9.

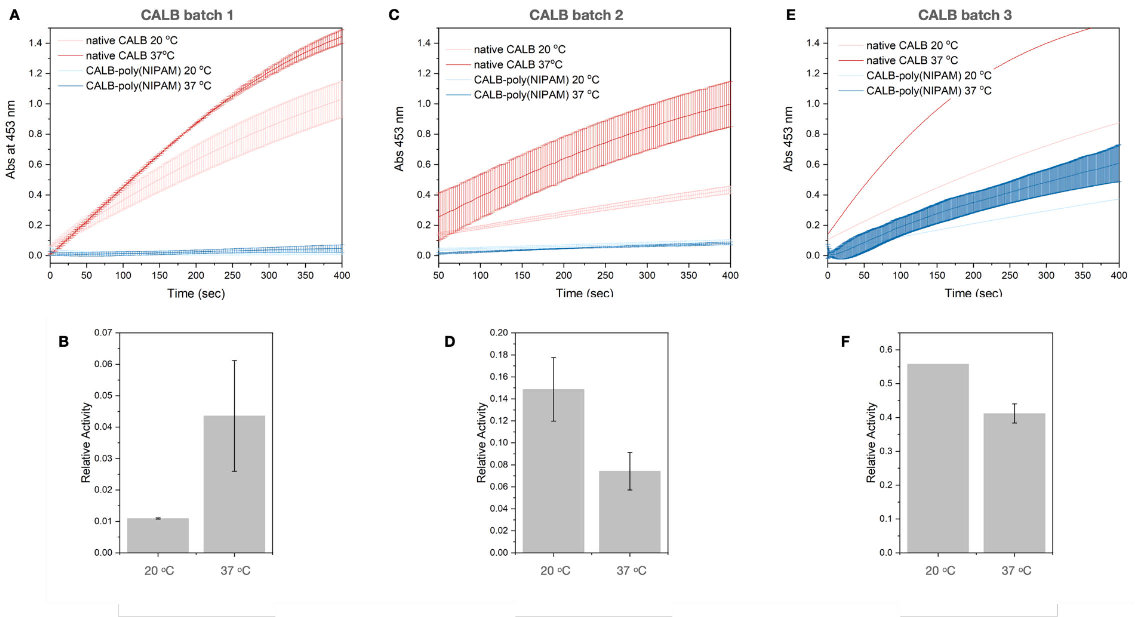


**Figure S39. A**. Esterase-like activity of batch 1 native CALB and CALB-poly(NIPAM) at 20 and 37 °C on CFDA substrate. **B**. Relative activity of batch 1 CALB-poly(NIPAM) to native CALB at each temperature. **C**. Esterase-like activity of batch

2 native CALB and CALB-poly(NIPAM) at 20 and 37 $^{o}$C on CFDA substrate. **D**. Relative activity of batch 2 CALB-poly(NIPAM) to native CALB at each temperature. **E**. Esterase-like activity of batch 3 native CALB and CALB-poly(NIPAM) at 20 and 37 $^{o}$C on CFDA substrate. **F**. Relative activity of batch 3 CALB-poly(NIPAM) to native CALB at each temperature.
